# CXCR6⁺ natural killer cell immunotherapy preserves CD4⁺ T helper cells in humanized mice

**DOI:** 10.64898/2026.05.28.728486

**Authors:** Naushad Khan, Kayla Frank, Silke Paust

**Author notes:** **Corresponding author:** Silke Paust, Ph.D., The Jackson Laboratory for Genomic Medicine, 10 Discovery Road, Farmington, CT, USA.

## Abstract

Human immunodeficiency virus (HIV) persists despite antiretroviral therapy because long-lived viral reservoirs are not eliminated, and ongoing or rebound infection contributes to progressive loss of CD4⁺ T helper cells. Natural killer (NK) cells can acquire adaptive, antigen-experienced functions, including recall responses to HIV envelope protein, suggesting that defined NK-cell subsets may be therapeutically useful against HIV. Because HIV-responsive adaptive NK-cell activity is enriched among CXC chemokine receptor 6-positive (CXCR6⁺) NK cells, we tested whether CXCR6⁺ NK cells provide enhanced antiviral activity and CD4⁺ T-cell protection compared with CXCR6⁻ NK cells. In co-cultures with HIV-infected primary CD4⁺ T cells, PBMC-derived CXCR6⁺ and CXCR6⁻ NK cells both reduced viral replication, but CXCR6⁺ NK cells mediated significantly greater suppression. In HIV-infected humanized mice, weekly infusion of expanded PBMC-derived NK cells lowered plasma viral burden, with CXCR6⁺ NK cells providing stronger preservation of circulating CD4⁺ T cells and significant preservation of splenic CD4⁺ T cells. HIV-Env vaccination further enriched NK cells with enhanced therapeutic activity. CXCR6⁺ NK cells derived from HIV-Env–vaccinated humanized mice produced the strongest suppression of HIV replication and restored CD4⁺ T-cell frequencies in blood and spleen to levels comparable to uninfected controls. Together, these findings identify CXCR6⁺ NK cells as an HIV-responsive adaptive NK-cell subset that combines antiviral activity with preservation of CD4⁺ T-cell immunity in vivo. These data support further development of CXCR6⁺ NK-cell therapy as a vaccine-informed cellular immunotherapy strategy for HIV.

## 1 Introduction

Over 38.4 million people currently live with human immunodeficiency virus (HIV)-1, and 650,00 HIV-1-related deaths were reported in 2021 ^1^. HIV remains a global health burden because lifelong antiretroviral therapy is required to suppress viremia, and viral persistence is maintained by long-lived latent reservoirs that are not eliminated by current treatment ^2^. Viral persistence is coupled to progressive immunologic injury, including depletion and dysfunction of CD4⁺ Th cells ^3, 4^, which is the central immunologic defect underlying AIDS ^5, 6^. Even with durable viral suppression, chronic immune activation ^7^ and incomplete immune reconstitution contribute to co-infections ^8, 9^ and morbidity ^10^. Thus, there is strong interest in immunotherapies that not only suppress HIV replication but also preserve or restore CD4⁺ Th-cell homeostasis.

NK cells are cytotoxic innate lymphocytes that restrict viral infection and malignancy through perforin-and granzyme-mediated killing, death receptor pathways, and cytokine and chemokine secretion ^11^. Their activity is governed by integrated signaling from inhibitory and activating receptors, including killer immunoglobulin-like receptors, CD94/NKG2 family receptors such as NKG2A and NKG2D, and natural cytotoxicity receptors, enabling rapid responses to infected or stressed target cells ^11^. In HIV infection, NK cells contribute to immune pressure through direct cytotoxicity, antibody-dependent effector function, and immunoregulatory crosstalk that shapes adaptive immunity ^12–22^.

A key shift in the field is the recognition that NK cells can display adaptive features classically attributed to T and B lymphocytes, including antigen-specific recall responses^23–26^. A particularly well-supported axis of NK-cell memory involves CXCL16-dependent, liver-associated NK cells that express the chemokine receptor CXCR6^23, 24^. In mice, hepatic NK cells develop antigen-specific memory to structurally diverse antigens, including viral antigens such as influenza and HIV-1 vaccine antigens, and depend on the CXCR6/CXCL16 axis for longevity ^24^. Similarly, antigen-specific NK-cell memory to SIV and SHIV antigens has been demonstrated in nonhuman primates after infection or vaccination ^25^. In humanized mouse systems, vaccination with HIV envelope antigens can prime human NK cells, particularly CXCR6^+^ liver-resident NK populations, for vaccination-dependent, antigen-specific recall cytotoxicity ^23^. More recent studies have further refined this concept by demonstrating antigen-specific human NK-cell responses to HIV and influenza peptides, including responses linked to the NKG2/HLA-E axis ^26^. Together, these studies support the idea that CXCR6⁺ NK cells may represent an adaptive, antigen-experienced NK-cell population that can be therapeutically harnessed against HIV.

Multiple lines of evidence support NK cells as a viable therapeutic platform in HIV: NK cells can suppress HIV replication ex vivo, can be expanded and engineered, and can be adoptively transferred in vivo ^11, 27^. Importantly, next-generation humanized mouse studies have provided direct evidence that human NK cells can restrict HIV-1 infection and influence viral burden, supporting NK-cell therapy as a potential complement to antiretroviral and antibody-based strategies ^21, 28–30^. However, a clinically consequential gap remains: most NK-focused HIV therapeutic studies have prioritized virologic control^21^, whereas the central driver of HIV-associated immunodeficiency is the progressive loss and dysfunction of CD4⁺ Th cells ^5, 6^. Durable preservation of CD4⁺ T cells is essential for immune reconstitution, vaccine responsiveness, and resistance to opportunistic infection ^7–10^. Therefore, the critical translational question is not only whether an NK-cell product can reduce viral burden, but whether it can simultaneously protect CD4⁺ T-cell homeostasis in vivo.

Here, we tested the hypothesis that CXCR6⁺ NK cells represent an HIV-responsive adaptive NK-cell subset with enhanced antiviral function and a unique capacity to preserve CD4⁺ Th cells during infection. Using human donor PBMC-derived CXCR6⁺ and CXCR6⁻ NK cells expanded ex vivo, a humanized mouse HIV infection model with weekly adoptive NK-cell transfer, and HIV-Env vaccination to enrich and program humanized mouse–derived liver NK cells before therapeutic transfer, we define a CXCR6-linked NK-cell program that couples viral control with CD4⁺ T-cell preservation. This dual effect directly addresses the central immunopathology of HIV and supports CXCR6⁺ NK cells as a candidate immunotherapeutic platform for HIV.

## Materials and Methods

### Animal procedures and biosafety approval.s

All animal procedures were approved by The University of Connecticut Health Science Center Institutional Animal Care and Use Committee (UCHC IACUC) and were performed in accordance with the National Institutes of Health Guide for the Care and Use of Laboratory Animals. In vitro HIV experiments were performed under approved BSL-2 and A-BSL-2 biosafety approvals. Commercially obtained, de-identified human peripheral blood mononuclear cells (PBMCs) were used as a source of NK cells, and the project was reviewed by the Jackson Laboratorie’s Institutional Review Board and determined not to constitute human subjects research under 45 CFR 46.102(e).

### Humanized mice, criteria for reconstitution, and euthanasia

Female NOD.Cg-Prkdcscid Il2rgtm1Wjl Tg(IL15)1Sz/SzJ mice humanized with umbilical cord derived CD34^+^ hematopoietic stem cells were purchased from The Jackson Laboratory, Bar Harbor, ME (Stock No. 703089). Mice were included if human CD45^+^ cells comprised more than 60 percent of total CD45^+^ cells (human plus mouse). Mice not meeting these criteria were excluded. All experimental procedures were approved by the UConn Health Connecticut Institutional Animal Care and Use Committee. All mice appeared healthy as per a veterinary check, had not been previously used in experimentation, and were euthanized using an IACUC approved protocol as follows: Mice were anesthetized by inhalation of 3% isoflurane in oxygen, adequate anesthesia was confirmed by a firm bilateral foot pinch, followed by cervical dislocation performed by trained, IACUC approved personnel.

### HIV-Envelope glycoprotein immunization

Humanized NSG Tg(Hu IL15) mice (female, 17 to 23 weeks) were assigned to PBS control or HIV envelope vaccination groups. For HIV-Env vaccination, humanized mice received 10^6^ intravenous syngeneic B cells loaded with 20ug (10 ug i.v. and 10 ug i.p) recombinant HIV-Env on days 0 and 20, while control mice received PBS. Six weeks after the final immunization, humanized mice were euthanized and their livers collected for either NK cell isolation or immune phenotyping by flow cytometry.

### HIV challenge and treatment groups

Female NOD.Cg-Prkdcscid Il2rgtm1Wjl Tg(IL15)1Sz/SzJ mice were anesthetized with 3% isoflurane in oxygen delivered by inhalation before intraperitoneal infection with HIV-1 Q23.17 (10^5^ pfu infectious units in 200 microliters). One week after infection, mice received weekly infusions for seven weeks of PBS (control), CXCR6^+^ NK cells, or CXCR6^−^ NK cells (10^6^ cells per mouse). Blood was collected weekly for plasma viral load quantification, and blood and spleen were analyzed at endpoint for CD4^+^ Th cell frequencies by flow cytometry.

### Flow cytometry staining and acquisition

Cells were stained with antibody panels listed in **Supplementary Table 1** and analyzed by flow cytometry. Viability was assessed using LIVE/DEAD fixable dye with DAPI (Life Technologies). Data were acquired on a Cytek Aurora flow cytometer and analyzed using FlowJo.

### Human peripheral blood mononuclear cells and NK cell enrichment

Cryopreserved peripheral blood mononuclear cells from commercially obtained healthy donors were thawed, and NK cells were enriched by negative selection using the EasySep Human NK Cell Isolation Kit (17955, STEMCELL Technologies), washed, counted, and prepared for flow cytometry-based sorting. NK cells were defined as CD45^+^, CD3^−^, CD56^+^. Peripheral blood mononuclear cell derived NK cells were stained, including CD56, CD3, and human CD45, and sorted into CXCR6^+^ and CXCR6^−^ subsets using a BD FACSymphony S6 cell sorter. Sorted subsets were expanded ex vivo for 15 days using a protocol previously published by our laboratory ^31^.

### Liver leukocyte and NK isolation

Humanized mouse liver tissue was mechanically dissociated through a 40-micrometer nylon mesh to generate single cell suspensions. Immune cells were enriched by Ficoll Paque density gradient centrifugation, washed in PBS with 2% bovine serum, and then either expanded for further in vitro or in vivo use, or analyzed by flow cytometry.

### Sorting and expansion of humanized mouse derived liver CXCR6+ and CXCR6 negative NK cells

Following liver immune cell isolation from vaccinated mice, NK cells were enriched by negative selection (EasySep Human NK Cell Isolation Kit, 17955), washed in PBS with 2% fetal bovine serum, and prepared for flow cytometry-based sorting of CXCR6^+^ and CXCR6^-^ NK cells. Sorted subsets were expanded ex vivo for 15 days using a protocol previously published by our laboratory ^31^.

### CXCL16 feeder cell line, feeder cell inactivation and feeder composition

To support expansion of CXCR6^+^ NK cells in culture, the B lymphoblastoid cell line RPMI 8866 (Sigma Aldrich) was engineered to stably express CXCL16, reflecting the CXCR6 CXCL16 survival axis. RPMI 8866 cells were transduced with lentiviral particles (pLenti C mGFP, Origene) carrying CXCL16 transcript variant 1 (TV1, NM_022059) or variant 2 (TV2, NM_001100812). Transduced feeder cells were sorted for GFP expression on a FACSymphony S6, and CXCL16 expression and absence of CXCR6 on RPMI 8866 cells were confirmed by post transduction flow cytometry. Feeder populations were treated with mitomycin C (10 micrograms per milliliter) and subsequently irradiated using an X ray irradiator (X Rad320, Precision, UConn Health). Allogeneic cryopreserved peripheral blood mononuclear cells were used as additional feeder cells and combined at a 2:1 ratio (PBMC feeders to RPMI 8866 CXCL16 feeders) to promote proliferation of CXCR6^+^ and CXCR6^−^ NK cells.

### Ex vivo expansion of CXCR6^+^ and CXCR6^−^ NK cells

Sorted CXCR6^+^ and CXCR6^−^ NK cells were expanded for 15 days as published by us previously ^31^. The frequency of CXCR6^+^ NK cells in each donor sample guided seeding density, typically 100 to 1,000 cells per well. Cultures were maintained at 37 degrees Celsius with 5 percent carbon dioxide and refreshed every 2 to 3 days by adding fresh medium and removing residual feeder cells. Culture medium consisted of RPMI supplemented with phytohemagglutinin (PHA, 2 micrograms per milliliter), interleukin 2 (100 international units per milliliter, PeproTech), interleukin 15 (10 nanograms per milliliter, BioLegend), interleukin 15 receptor Fc chimera (20 nanograms per milliliter, R and D Systems), and 10 percent human AB serum (Sigma Aldrich). On day 14, expanded NK cells were analyzed by flow cytometry to confirm phenotype and purity and were used for functional assays or infused into humanized mice.

### Intravenous infusion of expanded NK cell subsets

Expanded NK cell subsets were washed, counted, and resuspended in sterile PBS for intravenous infusion. HIV-infected humanized mice received 10^6^ NK cells per infusion in a volume of 200 microliters, administered weekly for seven weeks.

### CD4+ T cell isolation for HIV suppression assays

Human CD4+ T cells were isolated from peripheral blood mononuclear cells by negative selection (CD4+ T cell negative selection kit, 17952, STEMCELL Technologies). Purified CD4+ T cells were rested overnight in R10 medium with interleukin 2 (200 units per milliliter, PeproTech 200 02) before activation and infection.

### HIV suppression assay

The NK cell based in vitro HIV suppression assay was adapted from a published T cell-based suppression assay ^32^. CD4^+^ T cells were activated using phytohemagglutinin (5 micrograms per milliliter) and CD3/CD28 beads (11161D, Life Technologies) according to the manufacturer’s instructions. Activation was verified 2- 4 days later by assessment of CD25 expression on CD3^+^CD4^+^ cells. Activated CD4^+^ T cells were infected with HIV-1 Q23.17 at multiplicity of infection 0.01. Three hours after infection, day 15 expanded peripheral blood mononuclear cell derived CXCR6^+^ or CXCR6^−^NK cells were added at a 1 to 1 effector to target ratio and co cultured for 3 days. HIV infection was quantified by intracellular HIV Gag staining using an anti HIV Gag antibody (Beckman 6604665).

### Quantification of plasma HIV-1 viral load by reverse transcription quantitative polymerase chain reaction

Peripheral blood was collected via retro orbital bleeding. After a 30-minute incubation at room temperature, sera were isolated by centrifugation at 1,000 times g for 15 minutes. Viral RNA was extracted using the QIAamp MinElute Virus Spin Kit (Qiagen). One step reverse transcription quantitative polymerase chain reaction was performed using qScript XLT 1 Step RT qPCR ToughMix (Quantabio) with Q23.17 targeting primers and probe: 5 prime CCTGTACCGTCAGCGTTATT 3 prime, 5 prime CAAAGAGAAGAGTGGTGGAGAG 3 prime, and 6 FAM 5 prime TGCTTCCTG ZEN CTGCTCCTAAGAACC 3 prime IABkFQ.

### Statistics

Statistical analyses were performed using GraphPad Prism. Data are shown as individual biological replicates with mean ± SEM unless otherwise stated. Statistical tests were two-sided unless otherwise indicated. For p-value–based analyses, p < 0.05 was considered statistically significant. For false-discovery-rate–corrected analyses, FDR-adjusted q < 0.05 was considered statistically significant.

For comparisons between two independent groups, two-tailed unpaired Student’s t tests were used when variances were comparable, and Welch’s t tests were used when unequal variance correction was appropriate. For nonparametric comparisons between human cohorts, including the frequency of NK cells with detectable CXCR6 transcript expression in people living with HIV and healthy donors, Mann–Whitney U tests were used.

For endpoint comparisons among three or more groups within a single experiment, including CD4⁺ T-cell frequencies in blood and spleen, one-way ANOVA was used followed by Tukey’s multiple-comparisons test. These analyses were performed separately within each in vivo experiment: the PBMC-derived NK-cell transfer experiment and the HIV-Env–vaccinated humanized mouse liver-derived NK-cell transfer experiment.

For Th:NK coculture assays, triplicate wells were treated as technical replicates and averaged within each NK donor for each experimental condition. The resulting donor-level means from five independent commercially sourced peripheral blood NK-cell donors were used as biological replicates. The same commercial CD4⁺ T-cell target donor was used across conditions; therefore, statistical inference reflects reproducibility across NK donor biological replicates and does not assess variability across CD4⁺ T-cell target donors. Donor-matched comparisons among HIV-only, CXCR6⁻ NK-cell-treated, and CXCR6⁺ NK-cell-treated conditions were analyzed using paired two-tailed tests with Holm correction for multiple comparisons. Where percent HIV suppression was analyzed, suppression was calculated for each NK donor as 100 × [(HIV-only − NK-treated condition) / HIV-only], using the matched HIV-only control for that donor’s assay set as the reference. Data from these assays are presented as mean ± SD.

For multiparameter NK-cell phenotyping analyses, marker frequencies were compared between paired CXCR6⁺ and CXCR6⁻ NK-cell populations using paired t tests with false-discovery-rate correction across markers. Vaccine-associated changes in marker expression within CXCR6⁺ or CXCR6⁻ NK-cell subsets were also corrected for multiple testing using FDR correction. FDR-adjusted q values < 0.05 were considered significant.

For longitudinal plasma viral-load analyses, integrated viral burden was quantified for each mouse by calculating the area under the curve from week 0 through week 7 using the trapezoidal method. AUC values were compared among groups by one-way ANOVA followed by Tukey’s multiple-comparisons test. Viral loads were also compared at individual weekly time points using one-way ANOVA followed by Tukey-adjusted post hoc pairwise comparisons. Thus, AUC p values represent cumulative viral burden across the full study period, whereas weekly p values represent treatment effects at individual time points.

For cross-experiment comparison of CD4⁺ T-cell preservation between PBMC-derived and HIV-Env–vaccinated humanized mouse liver-derived NK-cell treatments, CD4⁺ T-cell rescue was normalized within each experiment to its own PBS and HIV control means using the following formula:

CD4⁺ T-cell rescue = [(NK-treated value − HIV control mean) / (PBS control mean − HIV control mean)] × 100.

Normalized CD4⁺ T-cell rescue values were compared between PBMC-derived and HIV-Env–vaccinated humanized mouse liver-derived NK-cell groups using Welch’s t tests performed separately for each tissue. Blood CD4⁺ T-cell rescue values were compared only with blood CD4⁺ T-cell rescue values, and spleen CD4⁺ T-cell rescue values were compared only with spleen CD4⁺ T-cell rescue values. For analyses testing whether the CXCR6⁺-specific effect differed by NK-cell source, two-way ANOVA was used with NK-cell source and CXCR6 status as factors, and the interaction term was reported separately by tissue. These cross-experiment p values represent normalized rescue comparisons and are distinct from the raw within-experiment endpoint CD4⁺ T-cell comparisons.

Exact p values, adjusted p values, q values, or significance thresholds are reported in the Results, figure legends, and/or Supplementary Table 3, as applicable.

## Results

### In humans, HIV infection is associated with increased CXCR6 expression on PB-NK cells

We have previously demonstrated murine and human CXCR6^+^ NK cells to have immunological memory to HIV-Env^23, 24^. To determine whether HIV-infection alters the expression of CXCR6 on human peripheral blood (PB) NK cells, we compared the frequency of NK cells with detectable CXCR6 mRNA in PB from people living with HIV (PLWH) with that of healthy controls (**Supplemental Table 2**). The frequency of CXCR6^+^ was significantly increased in PLWH compared with healthy donors (**Fig. 1**). The frequency of CXCR6-mRNA^+^ NK cells was significantly higher in PLWH than in healthy donors, median 1.17, n = 75, versus 0.59, n = 139, respectively, Mann–Whitney U = 3498, two-tailed p < 0.0001. This infection associated enrichment supports the concept that CXCR6 marks an HIV responsive NK subset. Accordingly, we tested whether the CXCR6-defined NK cell subset differs in antiviral activity and in its ability to preserve CD4^+^ Th cells, whose loss leads to AIDS in PLWH.

**Figure 1.**
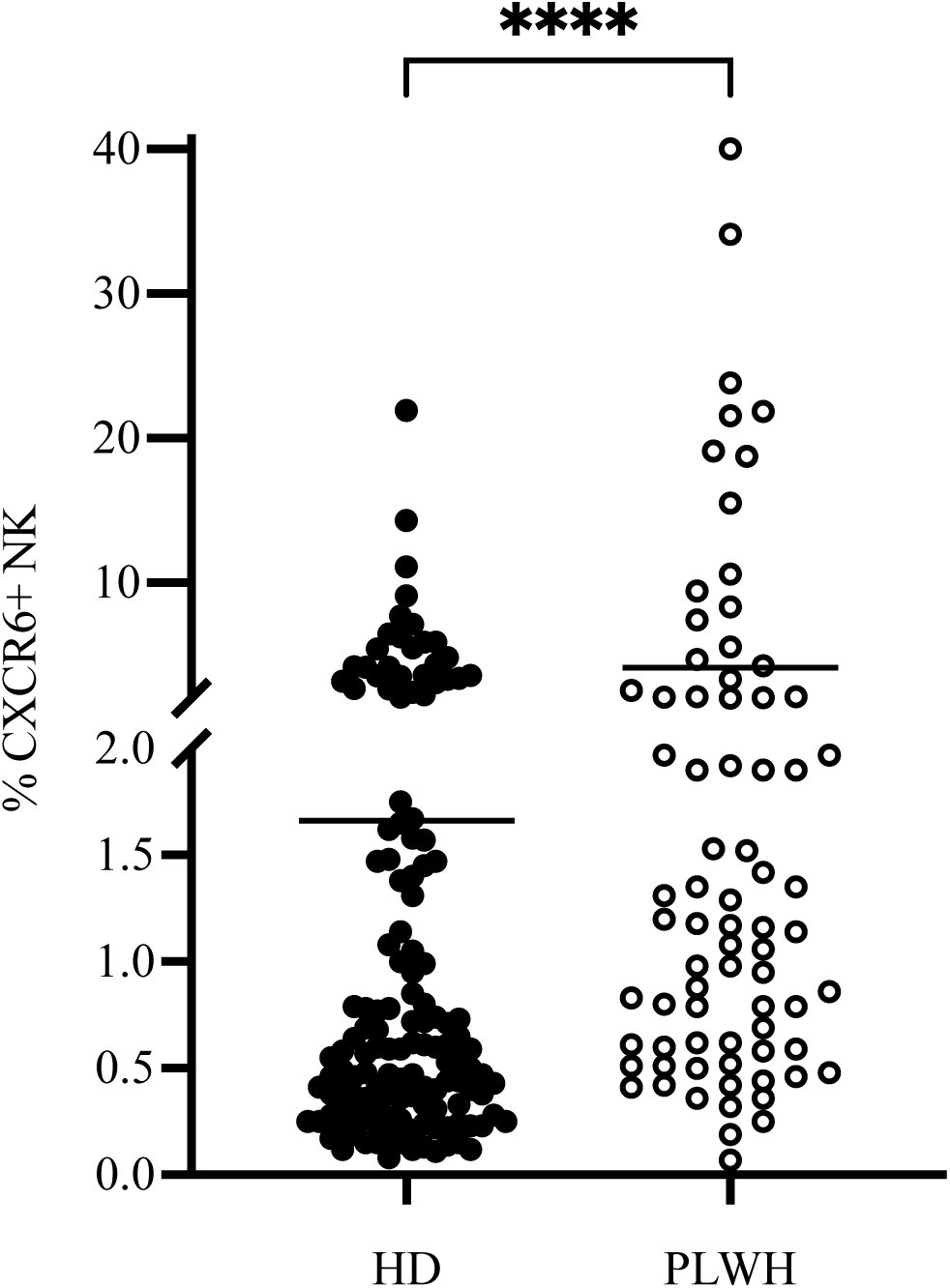
The frequency of CXCR6-mRNA-expressing NK cells is elevated in people living with HIV relative to healthy donors. Publicly available single-cell RNA-seq analysis was used to quantify the frequency of NK cells with detectable CXCR6 transcript expression in healthy donors (HD, n = 139) and people living with HIV (HIV+, n = 75). Each symbol represents one individual. Median frequencies were 1.17 in HIV+ individuals and 0.59 in HDs. CXCR6-mRNA+ NK-cell frequencies were significantly higher in HIV+ individuals than in HDs by two-sided Mann–Whitney U test (U = 3498, p < 0.0001). A detailed description of the publicly available datasets is included in the supplemental materials as **Supplemental Table 2**.

### CXCR6⁺ NK cells suppress HIV replication more effectively than CXCR6⁻ NK cells in a primary CD4⁺ T-cell co-culture assay

To enable functional comparison of CXCR6-defined NK cell subsets, we enriched NK cells from commercially obtained healthy donor PBMCs, sorted them into CXCR6⁺ and CXCR6⁻ fractions, and expanded them for 15 days using a protocol previously published by our laboratory ^31^. In brief, NK cells were stimulated with mitomycin C-treated, irradiated CXCL16-expressing RPMI 8866 feeder cells together with irradiated allogeneic PBMC feeders in RPMI supplemented with phytohemagglutinin, IL-2, IL-15, IL-15R Fc chimera, and 10% human AB serum. NK cells expanded approximately ten-fold over two weeks and remained >85% pure (**Fig. 2A, B**).

**Figure 2.**
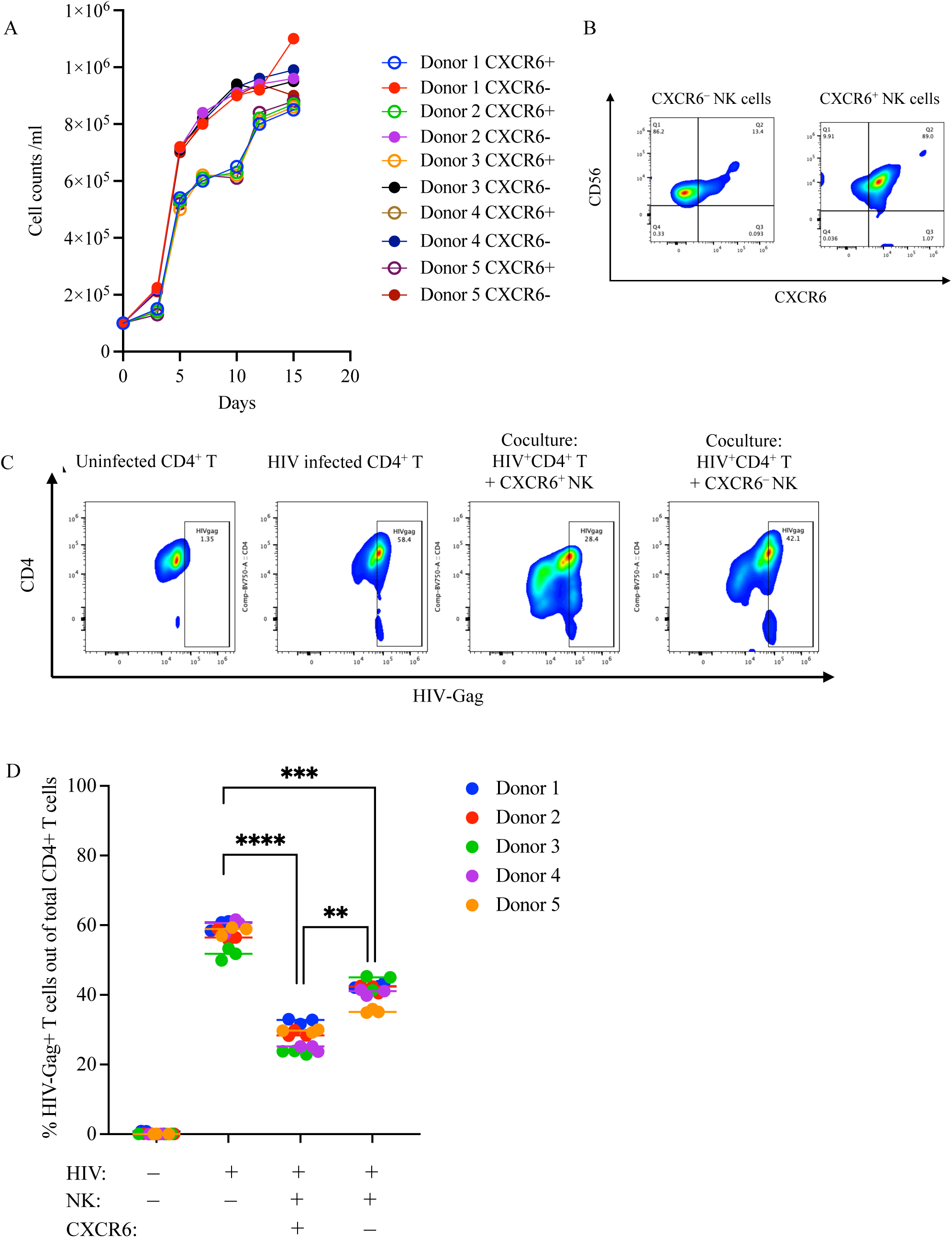
CXCR6⁺ NK cells suppress HIV replication more effectively than CXCR6⁻ NK cells in a primary CD4⁺ T-cell co-culture assay. A) Expansion kinetics of sorted CXCR6⁻ and CXCR6⁺ NK cells from independent peripheral blood NK-cell donors during 15 days of ex vivo culture with CXCL16-expressing feeder cells, irradiated allogeneic PBMC feeders, and cytokine supplementation. B) Representative post-expansion flow cytometry plots showing maintenance of CXCR6-defined NK-cell phenotypes following expansion. C) Representative intracellular HIV-Gag flow cytometry plots from the indicated culture conditions. Activated primary CD4⁺ T cells from a single peripheral blood donor were infected with HIV-1 Q23.17 at an MOI of 0.01 and cultured alone or co-cultured for 3 days with day 14-expanded CXCR6⁺ or CXCR6⁻ NK cells at a 1:1 effector-to-target ratio. HIV infection was quantified by intracellular HIV-Gag staining and flow cytometry. D) Quantification of HIV suppression assay results. CXCR6⁺ and CXCR6⁻ NK cells were tested using NK cells from five independent genetically unrelated donors, with the same HIV-infected CD4⁺ T-cell target donor used across all NK donor and subset conditions. For each NK donor and condition, the CD4^+^ T cell-frequency was determined by flow cytometry, triplicate technical wells per donor were averaged before statistical analysis, and the five NK donors were analyzed as biological replicates. Donor-level paired comparisons were analyzed using two-sided paired tests with Holm correction for multiple comparisons. CXCR6⁺ NK cells significantly reduced the frequency of HIV-Gag⁺ CD4⁺ T cells compared with HIV-only cultures, p = 0.000064. CXCR6⁻ NK cells also significantly reduced HIV-Gag⁺ CD4⁺ T cells compared with HIV-only cultures, p = 0.004059. CXCR6⁺ NK cells suppressed HIV significantly more effectively than paired CXCR6⁻ NK cells, p = 0.009637. Data are presented as mean ± SD, with each color representing an independent NK-cell donor. **p < 0.01, ****p < 0.0001.

We next evaluated expanded CXCR6⁺ and CXCR6⁻ effector NK cells from each of five genetically unrelated commercial peripheral blood NK donors in an in vitro HIV suppression assay ^33^, using activated, HIV-infected primary CD4⁺ T cells from a single commercial peripheral blood donor as targets. The same CD4⁺ T cell target donor was used across all NK donor and subset conditions to control for target-cell donor variability and enable direct comparison of NK subset-mediated suppression across independent NK donors. CD4⁺ T cell targets were activated and infected with HIV Q23.17 at a multiplicity of infection of 0.01, co-cultured with CXCR6⁺ or CXCR6⁻ NK cells at a 1:1 effector-to-target ratio, and infection was quantified after three days by intracellular HIV Gag staining and flow cytometric analysis (**Fig. 2C, D**). For each NK donor and condition, triplicate wells were performed and averaged before statistical analysis, with the five NK donors serving as biological replicates. In HIV-only cultures, the frequency of HIV-Gag⁺ CD4⁺ T cells was 57.43 ± 3.45%. CXCR6⁺ NK cells significantly reduced HIV-Gag⁺ CD4⁺ T cells to 27.83 ± 3.67%, corresponding to 51.59 ± 5.03% suppression relative to matched HIV-only controls (p = 0.000064, Holm-corrected). CXCR6⁻ NK cells also significantly suppressed HIV replication, reducing HIV-Gag⁺ CD4⁺ T cells to 40.90 ± 3.33%, corresponding to 28.44 ± 8.90% suppression (p = 0.004059, Holm-corrected). Importantly, CXCR6⁺ NK cells suppressed HIV significantly more effectively than paired CXCR6⁻ NK cells, with a mean difference of 23.15 percentage points in suppression (p = 0.009637, Holm-corrected).

Together, these data demonstrate that peripheral blood-derived CXCR6⁺ NK cells mediate potent suppression of HIV replication in vitro and are significantly more suppressive than paired CXCR6⁻ NK cells under controlled target-cell conditions, supporting further evaluation of CXCR6⁺ NK cells for future HIV immunotherapy design.

### Healthy donor PBMC-derived CXCR6⁺ NK cell infusion suppresses HIV replication in vivo and protects against HIV-mediated CD4⁺ Th cell depletion

We next tested whether CXCR6⁺ NK cells provide added benefit in vivo, with emphasis on preservation of CD4⁺ Th cells as a clinically consequential endpoint, alongside reduction of plasma viremia. Humanized NSG Tg(Hu IL15) mice were infected with HIV-1 Q23.17 and then treated weekly for seven weeks with PBS (controls), peripheral blood derived CXCR6⁺ NK cells, or peripheral blood derived CXCR6^−^ NK cells (10^6^ cells/mouse, n = 4 mice per group; **Supplemental Figures 1 and 2**). Weekly healthy donor-derived NK-cell treatments significantly reduced HIV replication compared with HIV-infected controls receiving no NK cells/PBS (**Fig. 3A, B**).

**Figure 3.**
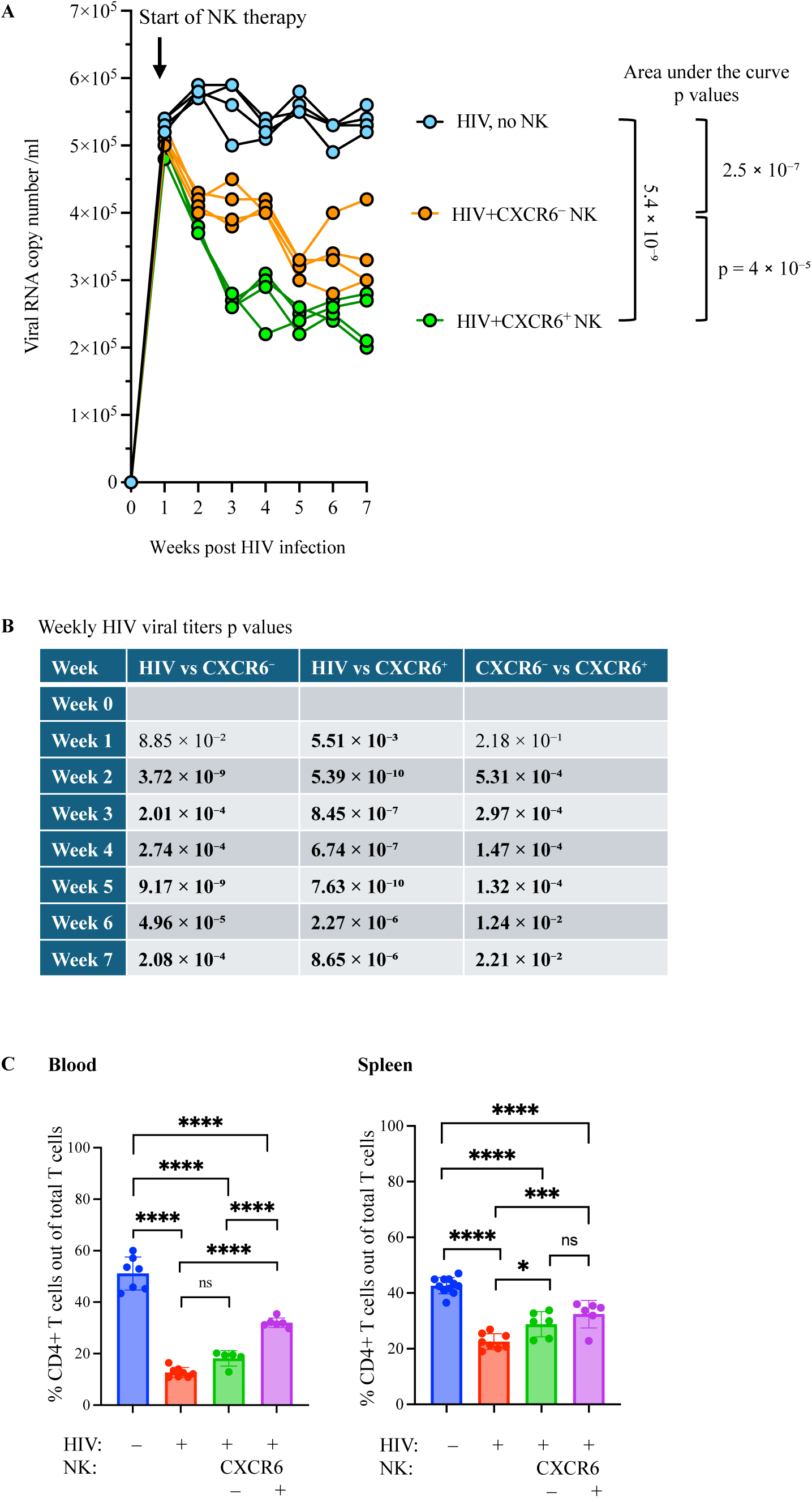
Healthy donor PBMC-derived CXCR6⁺ NK cell infusion suppresses HIV replication in vivo and protects against HIV-mediated CD4⁺ Th cell depletion. Humanized IL15TgNSG mice were intraperitoneally infected with HIV-1 Q23.17 followed by weekly intravenous infusion of PBS control, 10⁶ CXCR6⁺ NK cells, or 10⁶ CXCR6⁻ healthy donor PBMC-derived and expanded NK cells, with longitudinal weekly blood collection for assessment of viral load, and endpoint determination of viral load and CD4⁺ T-cell preservation in blood and spleen. **A**) Longitudinal plasma viral RNA analysis in Q23.17 HIV-1-infected humanized mice treated with PBS/no NK cells, 10⁶ CXCR6⁻ NK cells, or 10⁶ CXCR6⁺ NK cells. NK cell infusion reduced viral replication in vivo, with CXCR6⁺ NK cells producing the greatest suppression. Area under the curve analysis showed significant reductions in cumulative viral burden in CXCR6⁻ NK cell-treated mice and CXCR6⁺ NK cell-treated mice compared with HIV controls, with CXCR6⁺ NK cells significantly outperforming CXCR6⁻ NK cells. **B**) Tukey-adjusted post hoc comparisons of weekly viral titers. CXCR6⁺ NK cell treatment significantly reduced viral titers beginning at week 1, whereas CXCR6⁻ NK cell treatment became significant beginning at week 2. CXCR6⁺ NK cells showed significantly greater suppression than CXCR6⁻ NK cells from weeks 2 through 7. **C**) Quantification of CD4⁺ T cells as a percentage of total T cells in blood and spleen at endpoint. HIV infection significantly reduced CD4⁺ T-cell frequencies compared with HIV-negative PBS controls in both blood and spleen. In blood, CXCR6⁺ NK-cell treatment significantly preserved CD4⁺ T cells compared with HIV/no NK, whereas CXCR6⁻ NK-cell treatment did not. Direct comparison of NK-cell treatment groups showed significantly greater preservation of blood CD4⁺ T cells in the CXCR6⁺ NK-cell group compared with the CXCR6⁻ NK-cell group. In spleen, both CXCR6⁻ and CXCR6⁺ NK-cell treatment significantly increased CD4⁺ T-cell frequencies compared with HIV/no NK; however, the difference between CXCR6⁻ and CXCR6⁺ NK-cell treatment was not significant. Data are presented as mean ± SEM, with each point representing an individual mouse. Statistical analysis for panel C was performed using one-way ANOVA followed by Tukey’s multiple comparisons test. For panel C, exact Tukey-adjusted p values were as follows: in blood, PBS versus HIV/no NK, p = 1.67 × 10⁻¹⁴; PBS versus HIV + CXCR6⁻ NK, p = 4.73 × 10⁻¹²; PBS versus HIV + CXCR6⁺ NK, p = 5.16 × 10⁻⁸; HIV/no NK versus HIV + CXCR6⁻ NK, p = 0.0850; HIV/no NK versus HIV + CXCR6⁺ NK, p = 2.51 × 10⁻⁸; and HIV + CXCR6⁻ NK versus HIV + CXCR6⁺ NK, p = 3.16 × 10⁻⁵. In spleen, PBS versus HIV/no NK, p = 2.25 × 10⁻¹¹; PBS versus HIV + CXCR6⁻ NK, p = 3.09 × 10⁻⁷; PBS versus HIV + CXCR6⁺ NK, p = 4.66 × 10⁻⁵; HIV/no NK versus HIV + CXCR6⁻ NK, p = 0.0198; HIV/no NK versus HIV + CXCR6⁺ NK, p = 0.000182; and HIV + CXCR6⁻ NK versus HIV + CXCR6⁺ NK, p = 0.337. *p < 0.05, **p < 0.01, ***p < 0.001, ****p < 0.0001; ns, not significant.

Area under the curve (AUC) analysis demonstrated a significant effect of treatment group on total viral burden across the 7-week study period (p < 0.0001; **Fig. 3A**). AUC values were highest in HIV mice, with a mean ± SEM of 3,546,254 ± 32,875, followed by HIV^+^CXCR6⁻ NK cell-treated mice at 2,571,255 ± 64,884, and HIV+CXCR6⁺ NK cell-treated mice at 2,037,505 ± 27,727. Tukey post hoc analysis confirmed significant reductions in cumulative viral burden following NK cell infusion, with HIV vs HIV+CXCR6⁻ NK cells, p = 2.50 × 10⁻⁷; HIV vs HIV+CXCR6⁺ NK cells, p = 5.38 × 10⁻⁹; and HIV+CXCR6⁻ NK cells vs HIV+CXCR6⁺ NK cells, p = 3.99 × 10⁻⁵. Relative to HIV alone, cumulative viral burden was reduced by approximately 27.5% following infusion of CXCR6⁻ NK cells and by 42.5% following infusion of CXCR6⁺ NK cells. These findings demonstrate that adoptively infused NK cells significantly suppress HIV replication in vivo, with CXCR6⁺ NK cells mediating the strongest reduction in viral titers and cumulative viral burden (**Fig. 3A**).

Weekly time point analysis further demonstrated sustained antiviral activity of the infused NK cell populations (**Fig. 3B**). One-way ANOVA analysis revealed significant treatment effects at each weekly time point, with NK cell-treated groups exhibiting reduced viral titers compared with HIV alone. CXCR6⁺ NK cell-treated mice showed significantly lower viral titers than HIV mice beginning at week 1, whereas CXCR6⁻ NK cell-treated mice showed significant reductions beginning at week 2. From weeks 2 through 7, both NK cell-treated groups maintained significantly lower viral titers than HIV controls. Direct comparison of the NK cell subsets further showed that CXCR6⁺ NK cells produced significantly greater viral suppression than CXCR6⁻ NK cells from weeks 2 through 7 (**Fig. 3A, B**). Together, these longitudinal analyses indicate that infused NK cells exert durable suppression of viral replication in vivo, with the CXCR6⁺ NK subset displaying superior and sustained antiviral efficacy.

At endpoint, we quantified CD4⁺ Th cells in blood and spleen because CD4⁺ Th cells are primary targets of HIV and their progressive depletion causes AIDS. As expected, HIV infection induced marked depletion of CD4⁺ Th cells in both blood and spleen (**Fig. 3C**). In blood, CD4⁺ Th-cell frequency was significantly reduced from 51.14 ± 2.42% in HIV-negative PBS controls to 12.64 ± 0.69% in HIV-infected mice receiving no NK-cell therapy, p = 1.67 × 10⁻¹⁴. One-way ANOVA showed a significant overall treatment effect in blood, p = 1.80 × 10⁻¹⁴. CXCR6⁺ NK-cell infusion significantly protected against HIV-mediated Th-cell depletion, increasing blood Th-cell frequency to 32.00 ± 0.76% compared with HIV alone, p = 2.51 × 10⁻⁸. In contrast, CXCR6⁻ NK cell-treated mice showed only a modest increase to 18.16 ± 1.33%, which did not reach statistical significance compared with HIV alone, p = 0.0850. Direct comparison of the NK-cell subsets showed significantly greater preservation of blood CD4⁺ Th cells in the CXCR6⁺ NK cell-treated group compared with the CXCR6⁻ NK cell-treated group, p = 3.16 × 10⁻⁵, demonstrating superior CD4⁺ Th-cell protection by CXCR6⁺ NK cells in blood.

In the spleen, HIV infection also significantly reduced CD4⁺ Th-cell frequency from 42.65 ± 0.88% in HIV-negative PBS controls to 22.55 ± 1.00% in HIV-infected mice receiving no NK-cell therapy, p = 2.25 × 10⁻¹¹. One-way ANOVA showed a significant overall treatment effect in spleen, p = 4.15 × 10⁻¹¹. CXCR6⁻ NK cell-treated mice showed partial restoration of splenic CD4⁺ Th cells to 28.78 ± 1.85%, which reached statistical significance compared with HIV alone, p = 0.0198. CXCR6⁺ NK-cell infusion also significantly protected against HIV-mediated splenic CD4⁺ Th-cell loss, increasing splenic Th-cell frequency to 32.42 ± 2.01% compared with HIV alone, p = 0.000182. However, direct comparison of the NK-cell subsets showed no significant difference between CXCR6⁺ and CXCR6⁻ NK cell-treated groups in the spleen, p = 0.337. Together, these data indicate that infused NK cells protect against HIV-induced Th-cell depletion in both blood and spleen, with CXCR6⁺ NK cells showing the strongest protective effect in blood and both NK-cell subsets showing partial restoration in the spleen.

### CXCR6^+^ liver NK cells are phenotypically distinct at baseline and selectively remodeled by HIV-Env vaccination

To assess the impact of HIV-Env vaccination on the phenotype of human liver NK cells, humanized IL-15TgNSG mice were intravenously immunized with syngeneic B cells loaded with recombinant HIV-Env on days 0 and 20. Control mice received PBS. Six weeks after the final immunization, mice were euthanized, and liver NK cells were isolated and analyzed by flow cytometry for immune phenotyping. At baseline, CXCR6^+^ and CXCR6^−^ NK cells were phenotypically distinct, with 16 of 23 markers significantly different after False Discovery Rate (FDR) correction (**Fig. 4A, B**). Compared with CXCR6^−^ NK cells, control CXCR6^+^ NK cells expressed significantly higher NKp46, CD69, KIR3DL1, IFNγ, LILRB1, CD107a, NKp30, T-bet, and NKG2D, while expressing significantly lower CD57, KIR2DL1, Granzyme B, CD16, DNAM-1, NTB-A, and 2B4, demonstrating that CXCR6 expression identifies a distinct NK-cell subset at baseline. Following vaccination with HIV-Env, the CXCR6^+^ NK-cell compartment showed clear phenotypic remodeling (**Fig. 4C**), with significantly increased expression of 2B4 (+6.74, q=0.048), CD69 (+17.96, q=0.048), NTB-A (+21.16, q=0.048), NKp30 (+12.00, q=0.048), and Ki67 (+2.55, q=0.048), together with significantly decreased LILRB1 (−21.20, q=0.048) and NKp46 (−16.14, q=0.048). These vaccine-associated changes suggest activation and proliferative remodeling of CXCR6^+^ NK cells, accompanied by altered expression of activating and regulatory receptors. In contrast, HIV-Env vaccination did not result in FDR-significant phenotypic changes in CXCR6⁻ NK cells (**Fig. 4D**), supporting the conclusion that the vaccine-induced phenotypic response was concentrated in the CXCR6⁺ NK-cell subset.

**Figure 4.**
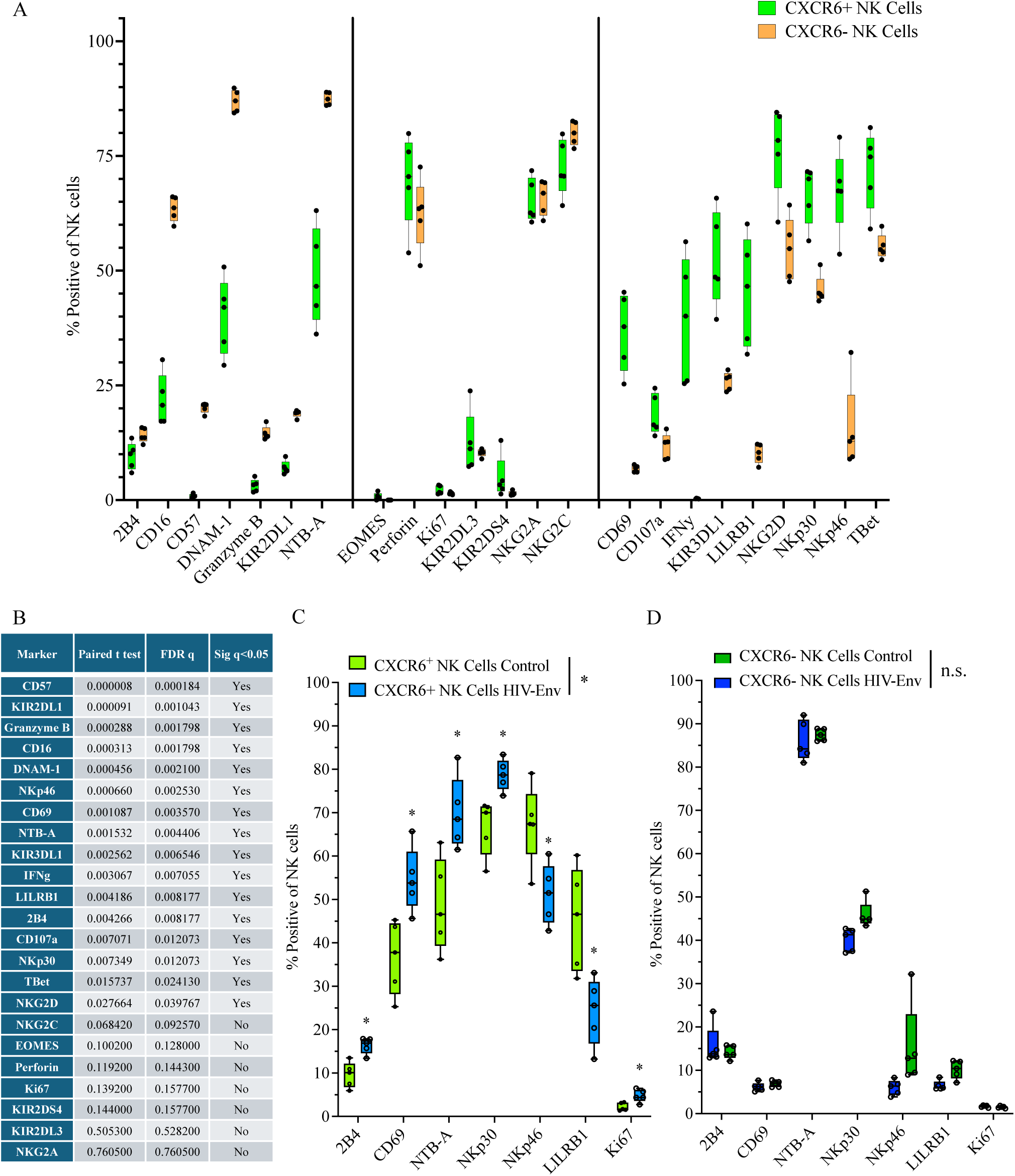
CXCR6^+^ liver NK cells are phenotypically distinct at baseline and selectively remodeled by HIV-Env vaccination. CD34^+^ humanized IL-15TgNSG mice were intravenously immunized with syngeneic B cells loaded with 20ug/mouse of recombinant HIV-Env on days 0 and 20, while control mice received PBS. Six weeks after the final immunization, mice were euthanized, and liver NK cells were phenotyped by flow cytometry. **A, B**) CXCR6^+^ and CXCR6^−^ NK cells from control mice showed distinct phenotypic profiles, with significant differences in CD57, KIR2DL1, granzyme B, CD16, DNAM-1, NKp46, CD69, NTB-A, KIR3DL1, IFNγ, LILRB1, 2B4, CD107a, NKp30, T-bet, and NKG2D after paired t test with false discovery rate (FDR) correction, q < 0.05; NKG2C, EOMES, perforin, Ki67, KIR2DS4, KIR2DL3, and NKG2A were not significant. **C**) HIV Env priming significantly altered the phenotype of CXCR6+ NK cells, including changes in 2B4, CD69, NTB-A, NKp30, NKp46, LILRB1, and Ki67, with q < 0.05 for each marker, whereas **D**) no significant changes were detected in CXCR6^−^ NK cells. Data are shown as percent marker-positive NK cells.

### HIV-Env–vaccinated humanized mouse liver-derived CXCR6⁺ NK cells suppress HIV replication in vivo and preserve CD4⁺ Th cells more effectively than CXCR6⁻ NK cells

Having established that PBMC-derived CXCR6⁺ NK cells suppress HIV replication in vivo, we next tested whether HIV-Env–vaccinated humanized mouse–derived liver NK cells showed comparable or enhanced antiviral activity. CXCR6⁺ and CXCR6^−^ NK cells were isolated from livers of HIV-Env vaccinated humanized mice and expanded using the same protocol used for expansion of PBMC-derived NK cells earlier (**Supplemental Figures 3 and 4**). Then, humanized NSG Tg(Hu IL15) mice were HIV-infected and treated weekly with PBS (control), HIV-Env–vaccinated humanized mouse–derived CXCR6^+^ or CXCR6^−^ NK cells, and plasma viral burden was monitored weekly (**Supplemental Figure 4**). Weekly HIV-Env–vaccinated humanized mouse–derived NK-cell treatments significantly reduced HIV replication compared with HIV-infected controls receiving no NK cells/PBS (**Fig. 5A, B**).

**Figure 5.**
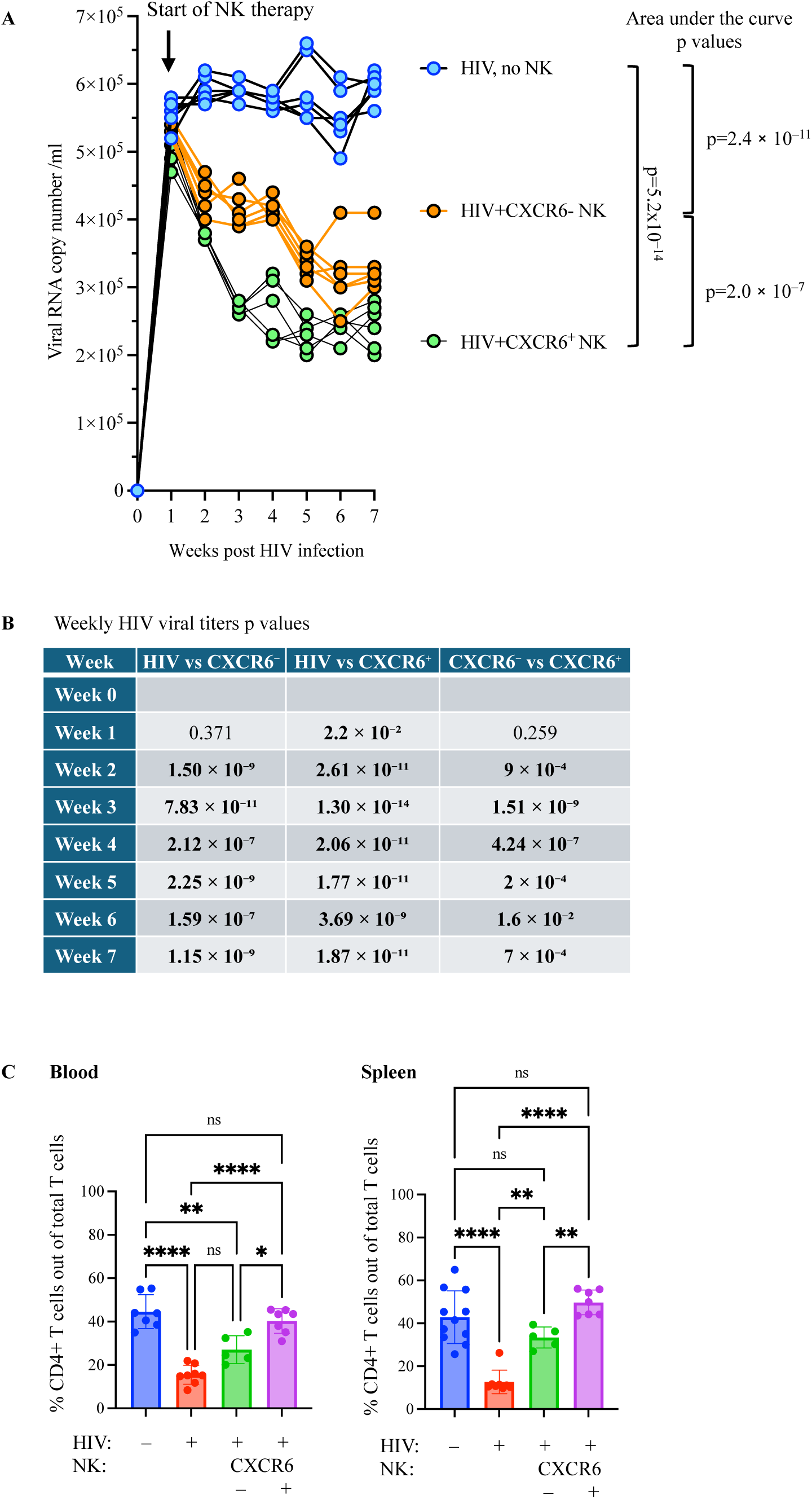
HIV-Env–vaccinated humanized mouse liver-derived CXCR6⁺ NK cells suppress HIV replication in vivo and preserve CD4⁺ Th cells more effectively than CXCR6⁻ NK cells. Humanized IL15TgNSG mice were intraperitoneally infected with HIV-1 Q23.17 followed by weekly intravenous infusion of PBS (control), 10^6^ CXCR6⁺, or 10^6^ CXCR6⁻ HIV-Env vaccinated humanized mouse derived liver NK cells, with longitudinal weekly blood collection for assessment of viral load, and endpoint determination of viral load and blood and spleen CD4^+^ Th cell preservation. **A)** Longitudinal plasma viral RNA analysis in Q23.17 HIV-1-infected humanized mice treated 10^6^ with no NK cells (control), 10^6^ CXCR6⁻ NK cells, or 10^6^ CXCR6⁺ NK cells derived from HIV-Env vaccinated humanized mice. NK cell infusion reduced viral replication in vivo, with CXCR6⁺ NK cells producing the greatest suppression. Area under the curve analysis showed significant reductions in cumulative viral burden in CXCR6⁻ NK cell-treated mice and CXCR6⁺ NK cell-treated mice compared with HIV controls, with CXCR6⁺ NK cells significantly outperforming CXCR6⁻ NK cells. **B)** Tukey-adjusted post hoc comparisons of weekly viral titers. CXCR6⁺ NK cell treatment significantly reduced viral titers beginning at week 1, whereas CXCR6⁻ NK cell treatment became significant beginning at week 2. CXCR6⁺ NK cells showed significantly greater suppression than CXCR6⁻ NK cells from weeks 2 through 7. **C)** Quantification of CD4⁺ T cells as a percentage of total T cells in blood and spleen. HIV infection significantly reduced CD4⁺ T cell frequencies compared with PBS controls. In blood, CXCR6⁺ NK cell treatment significantly preserved CD4⁺ T cells compared with HIV alone and provided greater protection than CXCR6⁻ NK cells. In spleen, both CXCR6⁻ and CXCR6⁺ NK cells significantly protected against HIV-induced CD4⁺ T cell depletion, with CXCR6⁺ NK cells producing significantly greater restoration than CXCR6⁻ NK cells. Data are presented as mean ± SEM, with each point representing an individual mouse. Statistical analysis was performed using one-way ANOVA followed by Tukey’s multiple comparisons test. *p < 0.05, **p < 0.01, ****p < 0.0001; ns, not significant.

Area-under-the-curve analysis showed that cumulative viral burden was reduced by HIV-Env–vaccinated humanized mouse–derived CXCR6⁻ NK cells, p = 2.4 × 10⁻¹¹, and more strongly by HIV-Env–vaccinated humanized mouse–derived CXCR6⁺ NK cells, p = 5.2 × 10⁻¹⁴ (**Fig. 5A**). Direct comparison of the two HIV-Env–vaccinated humanized mouse–derived NK-cell subsets showed that CXCR6⁺ NK cells suppressed cumulative viral burden more effectively than CXCR6⁻ NK cells, p = 2.0 × 10⁻⁷.

Weekly viral-load analysis further demonstrated sustained antiviral activity by HIV-Env–vaccinated humanized mouse–derived NK cells (**Fig. 5B**). HIV-Env–vaccinated humanized mouse–derived CXCR6⁺ NK-cell–treated mice showed significantly lower viral titers than HIV controls beginning at week 1, p = 0.022, and this suppression was maintained through week 7. HIV-Env–vaccinated humanized mouse–derived CXCR6⁻ NK cells showed delayed but significant viral suppression beginning at week 2, p = 1.50 × 10⁻⁹. HIV-Env–vaccinated humanized mouse–derived CXCR6⁺ NK cells produced significantly greater viral suppression than HIV-Env–vaccinated humanized mouse–derived CXCR6⁻ NK cells from weeks 2 through 7, including week 2, p = 0.00093; week 3, p = 1.51 × 10⁻⁹; week 4, p = 4.24 × 10⁻⁷; week 5, p = 0.0002; week 6, p = 0.016; and week 7, p = 0.0007.

At endpoint, CD4⁺ Th-cell preservation was assessed in blood and spleen first within the HIV-Env–vaccinated humanized mouse–derived NK-cell experiment using raw CD4⁺ T-cell frequencies (**Fig. 5C**). In blood, HIV-Env–vaccinated humanized mouse–derived CXCR6⁺ NK cells significantly increased CD4⁺ T cells compared with HIV controls, p = 1.1 × 10⁻⁶, and compared with HIV-Env–vaccinated humanized mouse–derived CXCR6⁻ NK cells, p = 0.0062. Importantly, CD4⁺ T-cell frequencies in the HIV-Env–vaccinated humanized mouse–derived CXCR6⁺ NK-cell group were not significantly different from PBS controls, p = 0.26, indicating near-complete preservation of circulating CD4⁺ T cells. In spleen (**Fig. 5C**), HIV-Env–vaccinated humanized mouse–derived CXCR6⁺ NK cells also significantly increased CD4⁺ T cells compared with HIV controls, p = 1.6 × 10⁻⁸, and compared with HIV-Env–vaccinated humanized mouse–derived CXCR6⁻ NK cells, p = 0.00040. As in blood, splenic CD4⁺ T-cell frequencies in the HIV-Env–vaccinated humanized mouse–derived CXCR6⁺ NK-cell group were not significantly different from PBS controls, p = 0.17. Thus, HIV-Env–vaccinated humanized mouse–derived CXCR6⁺ NK cells significantly protected CD4⁺ T cells and restored CD4⁺ T-cell frequencies to PBS-like levels in both blood and spleen.

To directly compare CD4⁺ T-cell preservation across the two independent in vivo experiments shown in **Figs. 3 and 5**, we performed a separate normalized CD4 rescue analysis. Because the PBMC-derived NK-cell transfer experiment and the HIV-Env–vaccinated humanized mouse–derived NK-cell transfer experiment had separate PBS and HIV control groups, CD4⁺ T-cell rescue was normalized within each experiment to its own control means using the formula: [(NK-treated group − HIV control mean)/(PBS control mean − HIV control mean)] × 100. These values therefore represent normalized cross-experiment rescue comparisons, not pooled raw endpoint CD4⁺ T-cell frequencies. This analysis is not plotted in **Fig. 5** and is summarized in **Supplementary Table 3**.

Using this normalized analysis, both PBMC-derived and HIV-Env–vaccinated humanized mouse–derived CXCR6⁺ NK cells protected against HIV-associated CD4⁺ T-cell loss; however, HIV-Env–vaccinated humanized mouse–derived CXCR6⁺ NK cells produced significantly stronger preservation. In blood, PBMC-derived CXCR6⁺ NK cells produced approximately 50% CD4 rescue, whereas HIV-Env–vaccinated humanized mouse–derived CXCR6⁺ NK cells produced approximately 85% rescue, p = 0.0027. In spleen, PBMC-derived CXCR6⁺ NK cells produced approximately 49% rescue, whereas HIV-Env–vaccinated humanized mouse–derived CXCR6⁺ NK cells produced approximately 122% rescue, p = 0.00018. When blood and spleen were analyzed together, HIV-Env–vaccinated humanized mouse–derived CXCR6⁺ NK cells showed significantly greater overall CD4⁺ T-cell rescue than PBMC-derived CXCR6⁺ NK cells, 104% versus 50% rescue, p = 2.6 × 10⁻⁶.

Within the PBMC-derived NK-cell experiment, CXCR6⁺ NK cells significantly protected against HIV-associated CD4⁺ T-cell loss, but CD4⁺ T-cell frequencies remained below PBS control levels, indicating partial preservation. In contrast, within the HIV-Env–vaccinated humanized mouse–derived NK-cell experiment, CXCR6⁺ NK cells restored CD4⁺ T-cell frequencies to levels that were not significantly different from PBS controls. Thus, both the within-experiment endpoint analysis and the normalized cross-experiment rescue analysis support the conclusion that HIV-Env–vaccinated humanized mouse–derived CXCR6⁺ NK cells mediate stronger CD4⁺ T-cell preservation than PBMC-derived CXCR6⁺ NK cells.

Notably, HIV-Env–vaccinated humanized mouse–derived CXCR6⁻ NK cells also showed greater normalized CD4⁺ T-cell preservation than PBMC-derived CXCR6⁻ NK cells, particularly in spleen. HIV-Env–vaccinated humanized mouse–derived CXCR6⁻ NK cells showed a trend toward greater blood CD4 rescue than PBMC-derived CXCR6⁻ NK cells, p = 0.059, and significantly greater splenic rescue, p = 0.011; combined analysis also favored HIV-Env–vaccinated humanized mouse–derived CXCR6⁻ NK cells, p = 0.0045. Therefore, part of the advantage associated with HIV-Env–vaccinated humanized mouse–derived NK cells may reflect broader effects of NK-cell source, **tissue origin,** or priming rather than CXCR6 expression alone. However, the CXCR6⁺-specific enhancement was greater in HIV-Env–vaccinated humanized mouse–derived NK cells than in PBMC-derived NK cells in spleen, p = 0.048, but not in blood, p = 0.49.

Together, these data indicate that HIV-Env vaccination in humanized mice enriches for NK cells with superior antiviral and immune-preserving activity, with the strongest effect observed in the CXCR6⁺ NK-cell subset.

## Discussion

This study identifies CXCR6⁺ NK cells, particularly adaptive HIV-Env-primed CXCR6⁺ NK cells, as a candidate cellular immunotherapy that couples two clinically important outcomes in HIV infection: suppression of viral replication and preservation of CD4⁺ Th cell immunity. In ex vivo suppression assays, PBMC-derived CXCR6⁺ NK cells reduced HIV Gag expression in infected CD4⁺ T-cell targets more effectively than paired CXCR6⁻ NK cells. This antiviral hierarchy was maintained in vivo, where PBMC-derived CXCR6⁺ NK-cell infusion reduced cumulative plasma viral burden and significantly protected blood and splenic CD4⁺ T cells from HIV-associated depletion. Importantly, adaptive HIV-Env-primed CXCR6⁺ NK cells produced an even stronger therapeutic effect, reducing cumulative viral burden more effectively than HIV-Env-vaccinated humanized mouse-derived CXCR6⁻ NK cells and restoring CD4⁺ T-cell frequencies to PBS-like levels in both blood and spleen. After normalization to experiment-specific PBS and HIV controls, HIV-Env–vaccinated humanized mouse–derived CXCR6⁺ NK cells significantly outperformed PBMC-derived CXCR6⁺ NK cells in blood, spleen, and combined CD4⁺ T-cell rescue analyses, supporting the conclusion that HIV-Env vaccination enriches for an NK-cell population with superior antiviral and immune-preserving activity.

Our phenotypic analysis further supports the postulate that CXCR6 expression identifies a biologically distinct and vaccine-responsive NK-cell subset. In control cohort 1, CXCR6+ NK cells differed significantly from CXCR6– NK cells across multiple markers, with higher CD69, NKp46, NKp30, CD107a, IFNγ, LILRB1, KIR3DL1, T-bet, and NKG2D, and lower CD16, CD57, Granzyme B, DNAM-1, NTB-A, and 2B4. This baseline distinction is consistent with prior work showing that human CXCR6+ peripheral blood NK cells have a distinct, less mature/tissue-associated phenotype compared with CXCR6– NK cells. Importantly, vaccination induced significant phenotypic remodeling primarily within the CXCR6^+^ NK-cell compartment, increasing 2B4/CD244, CD69, NTB-A, NKp30, and Ki67, while decreasing LILRB1 and NKp46; in contrast, CXCR6^−^ NK cells showed no FDR-significant vaccine-induced marker changes in cohort 1. Thus, although prior studies used in vitro cytokine-driven PBMC/NK-cell expansion whereas the present study used vaccination of humanized mice, both support the concept that CXCR6^+^ NK cells are phenotypically distinct at baseline and retain stimulus-induced plasticity.

These findings are important because durable HIV control requires more than reduction of plasma viremia. Antiretroviral therapy suppresses viral replication but does not eliminate the stable latent reservoir in resting memory CD4⁺ T cells, which remains a major barrier to cure. Moreover, CD4⁺ T-cell loss and chronic immune activation are central drivers of HIV pathogenesis; microbial translocation and systemic inflammation contribute to immune dysfunction even beyond direct viral cytopathic effects. The ability of CXCR6⁺ NK-cell therapy to reduce viral burden while preserving CD4⁺ T cells therefore address two linked but distinct components of HIV disease: active viral replication and collapse of Th-cell homeostasis.

A major advance of the present work is that it links this adaptive NK-cell biology to therapeutic efficacy in an HIV challenge model. Prior studies have shown that human NK cells can restrict HIV replication in vivo. For example, NK-cell depletion in HIV-infected humanized MISTRG-6-15 mice increased plasma and tissue viral RNA and accelerated CD4⁺ T-cell loss, providing direct evidence that human NK cells can control HIV infection in vivo ^21^. The current study builds on that foundation by showing that therapeutic activity is enhanced by selecting CXCR6⁺ NK cells and further improved when those cells are derived from HIV-Env–vaccinated humanized mice. Thus, rather than treating NK cells as a bulk effector population, these data support a subset-defined and antigen-experienced NK-cell therapy strategy.

The CD4⁺ T-cell preservation observed here is especially notable considering previous immune-based HIV interventions. Interleukin-2 therapy increased CD4⁺ T-cell counts in large clinical studies, but in ESPRIT and SILCAAT, these numerical gains did not translate into clinical benefit compared with antiretroviral therapy alone ^34^. This distinction is important: increasing CD4⁺ T-cell numbers by nonspecific cytokine-driven expansion may not restore protective immunity. In contrast, CXCR6⁺ NK-cell transfer preserved CD4⁺ T cells while also suppressing HIV replication, suggesting that CD4⁺ T-cell rescue may reflect protection from infection-associated loss rather than simple lymphocyte expansion. This dual effect may be more relevant for immune reconstitution than CD4⁺ T-cell increase alone.

Several mechanisms could explain why CXCR6⁺ NK cells, and especially adaptive HIV-Env–primed CXCR6⁺ NK cells, preserve CD4⁺ T cells more effectively. First, enhanced killing of HIV-infected CD4⁺ T cells could reduce ongoing rounds of infection and thereby limit direct viral cytopathicity. Second, CXCR6-dependent trafficking or retention through CXCL16-rich tissue niches may improve localization to sites where HIV replication, immune activation, and CD4⁺ T-cell depletion are concentrated. Third, HIV-Env vaccination may imprint CXCR6⁺ NK cells with enhanced effector receptor programs, enabling more rapid recognition of infected targets. Fourth, CXCR6⁺ NK cells may indirectly preserve CD4⁺ T cells by modulating inflammatory tone, reducing bystander activation, or limiting tissue damage. This possibility is consistent with observations from natural SIV hosts, in which high viremia can coexist with preserved T-cell homeostasis and limited disease progression when immune activation is restrained. However, further studies are needed to evaluate these hypotheses.

The comparison between PBMC-derived and HIV-Env-vaccinated humanized mouse-derived NK cells also provides a useful translational insight. PBMC-derived CXCR6⁺ NK cells were clearly active, suppressing HIV replication and partially preserving CD4⁺ T cells. However, HIV-Env–vaccinated humanized mouse-derived CXCR6⁺ NK cells produced near-complete or complete CD4⁺ T-cell restoration relative to PBS controls. At the same time, HIV-Env-vaccinated humanized mouse–derived CXCR6⁻ NK cells also showed greater CD4⁺ T-cell preservation than PBMC-derived CXCR6⁻ NK cells, indicating that part of the advantage may reflect broader effects of NK-cell source, tissue origin, or HIV-Env vaccination rather than CXCR6 expression alone. Nonetheless, the CXCR6⁺-specific enhancement was strongest in spleen, where HIV replication is most pronounced in humanized mice, and where the interaction analysis supported a greater CXCR6⁺ advantage for HIV-Env–vaccinated humanized mouse–derived NK cells than for PBMC-derived NK cells.

Several limitations should be addressed in future work. The in vivo studies were performed in humanized mice, which are powerful for testing human immune cells during HIV infection but do not fully reproduce the chronic inflammatory, anatomical reservoir, and immune-regulatory complexity of human HIV disease. The comparison between PBMC-derived and HIV-Env–vaccinated humanized mouse–derived NK cells also required normalization across separate experimental settings, and the improved activity of HIV-Env–vaccinated humanized mouse–derived CXCR6⁻ NK cells indicates that CXCR6 expression, tissue source, and vaccine exposure should be disentangled experimentally. Future studies should track transferred NK cells in blood, spleen, lymph node, gut, and liver; define their transcriptional and epigenetic programs; test whether CXCR6 blockade or deletion reduces protection; and determine whether these cells reduce cell-associated HIV DNA or rebound after antiretroviral interruption.

In summary, our data support a model in which CXCR6⁺ NK cells represent an HIV-responsive adaptive NK-cell subset with therapeutic potential. PBMC-derived CXCR6⁺ NK cells suppress HIV replication and partially preserve CD4⁺ T cells, whereas HIV-Env–vaccinated humanized mouse–derived CXCR6⁺ NK cells show stronger and more complete immune preservation. By combining antigen experience, CXCR6-based subset selection, and adoptive transfer, this approach advances NK-cell therapy for HIV beyond nonspecific cytotoxicity and toward a vaccine-informed cellular product capable of both antiviral control and protection of CD4⁺ Th cell immunity.

## Author Contributions (CRediT)

Conceptualization: S.P.

Formal analysis: N.K., S.P.

Investigation: N.K., K.F.

Methodology: N.K., K.F., S.P.

Resources: S.P.

Supervision: S.P.

Visualization: N.K., S.P.

Writing original draft: N.K., S.P.

Writing review & editing: all authors

## Acknowledgments

We extend our appreciation to Blaine Pattavina and Himani Sharma for dedicated animal care throughout the research project, and we thank Himani Sharma for developing the flow cytometry panels used for these studies. We also thank the Flow Cytometry Core staff at Scripps Research and The Jackson Laboratory for Genomic Medicine for their technical expertise and support. We thank Tess Comeaux for careful proofreading.

## 11 Data Availability Statement

The datasets generated for this study can be obtained from the corresponding author by request.

## Supplementary Material

**Supplemental Figure 1.**
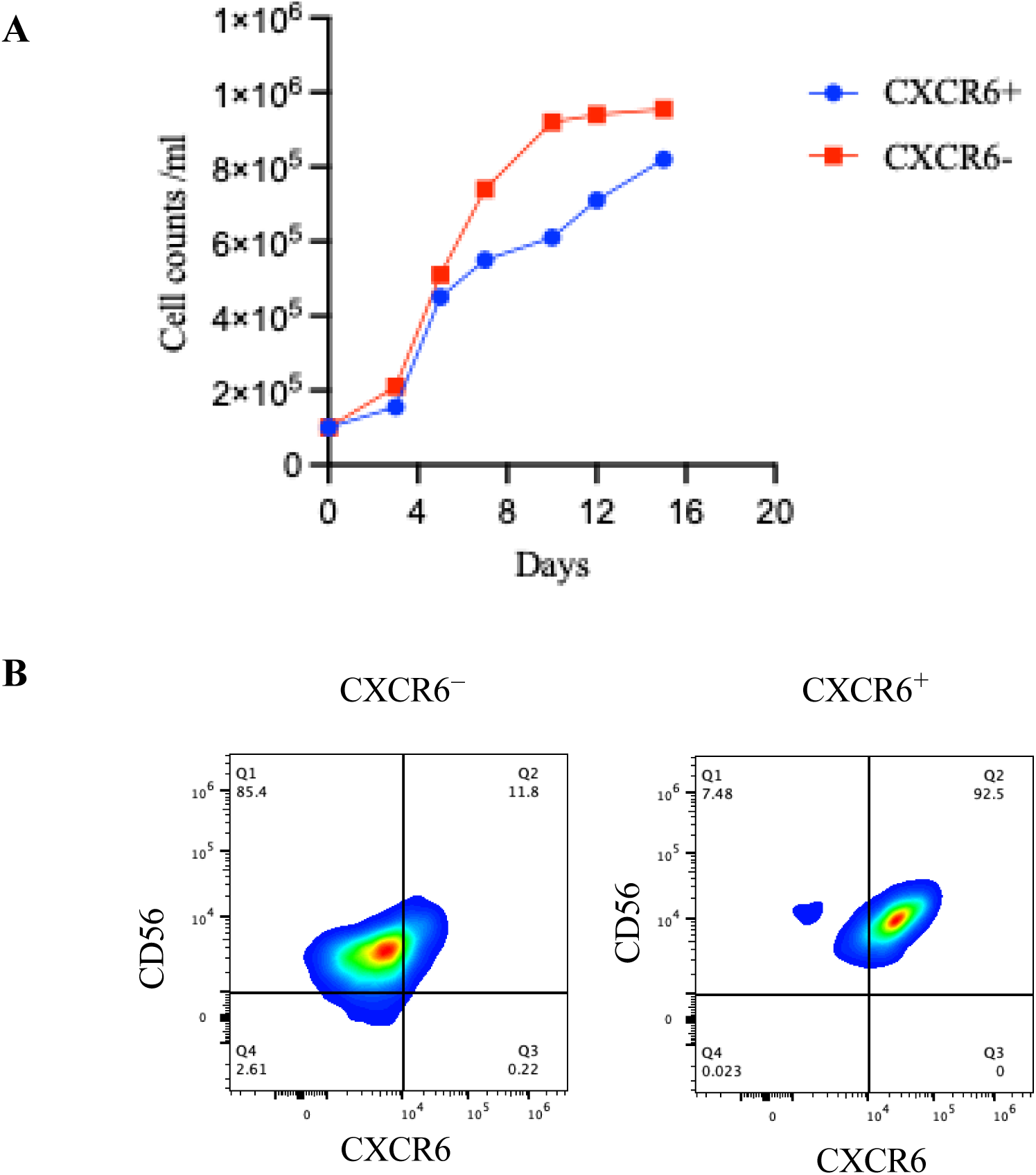
In vitro expansion and post-expansion purity of PBMC-derived CXCR6⁺ and CXCR6⁻ NK cells were assessed by cell counting and flow cytometry. **A)** Growth kinetics of expanded CXCR6⁺ and CXCR6⁻ NK cells are shown as cell counts per milliliter over 15 days. **B)** Representative flow cytometry plots show CD56 and CXCR6 expression in expanded CXCR6⁻ and CXCR6⁺ NK cell populations as determined by flow cytometry.

**Supplemental Figure 2.**
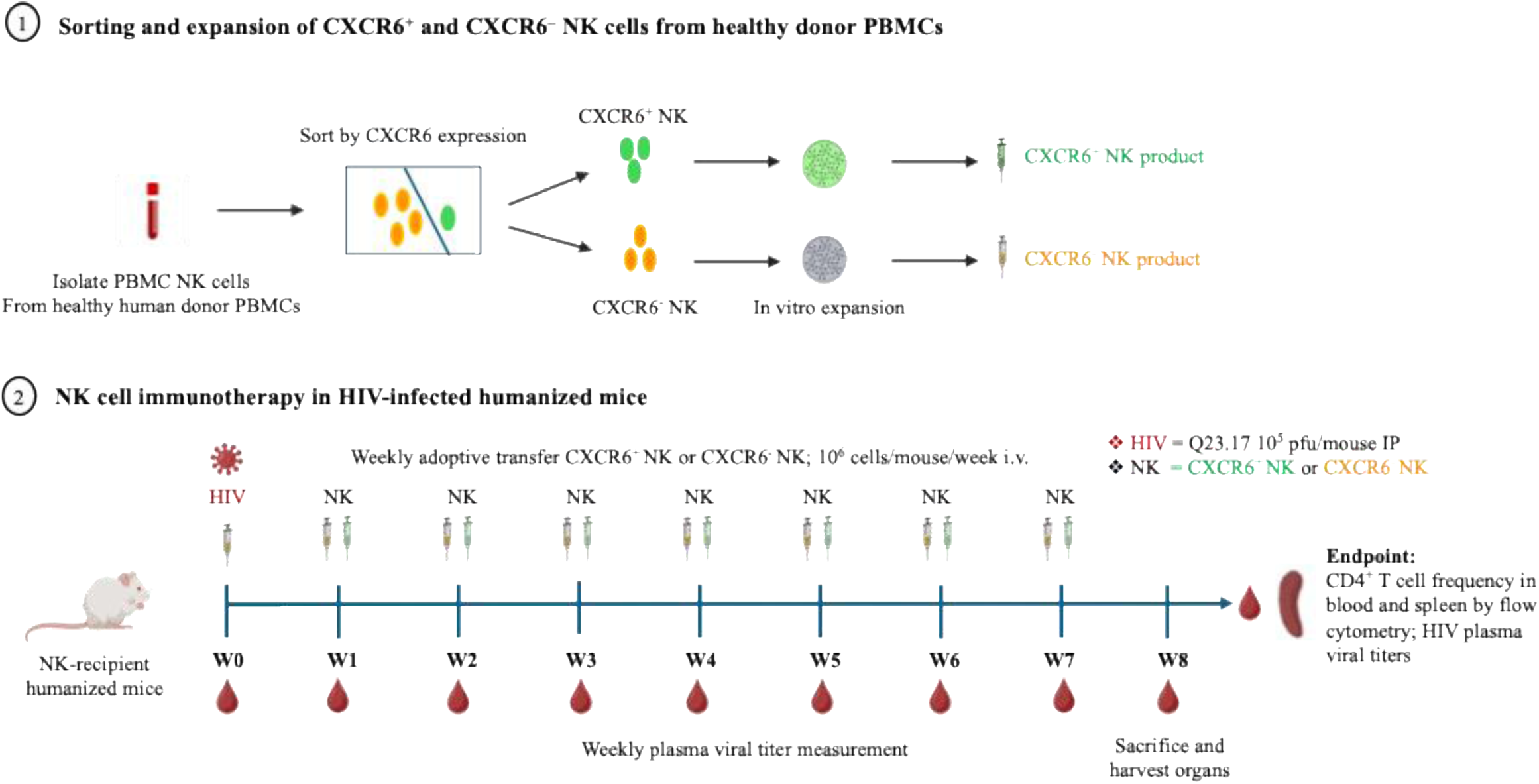
Experimental outline for NK cell immunotherapy using healthy donor PBMC derived NK cells. Healthy donor PBMC derived NK cells were sorted based on CXCR6 expression and expanded in vitro to generate CXCR6^+^ and CXCR6^−^ NK cell products. Umbilical cord blood-derived CD34^+^ HSC-humanized IL-15TgNSG mice were infected intraperitoneally with HIV-1 Q23.17, followed by weekly intravenous adoptive transfer of PBS control, 10⁶ CXCR6^+^ NK cells, or 10⁶ CXCR6^−^ NK cells per mouse. Plasma was collected longitudinally each week to measure HIV viral burden, and mice were sacrificed at week 8 for endpoint analysis of CD4^+^ T cell preservation in blood and spleen by flow cytometry, together with assessment of plasma viral titers.

**Supplemental Figure 3.**
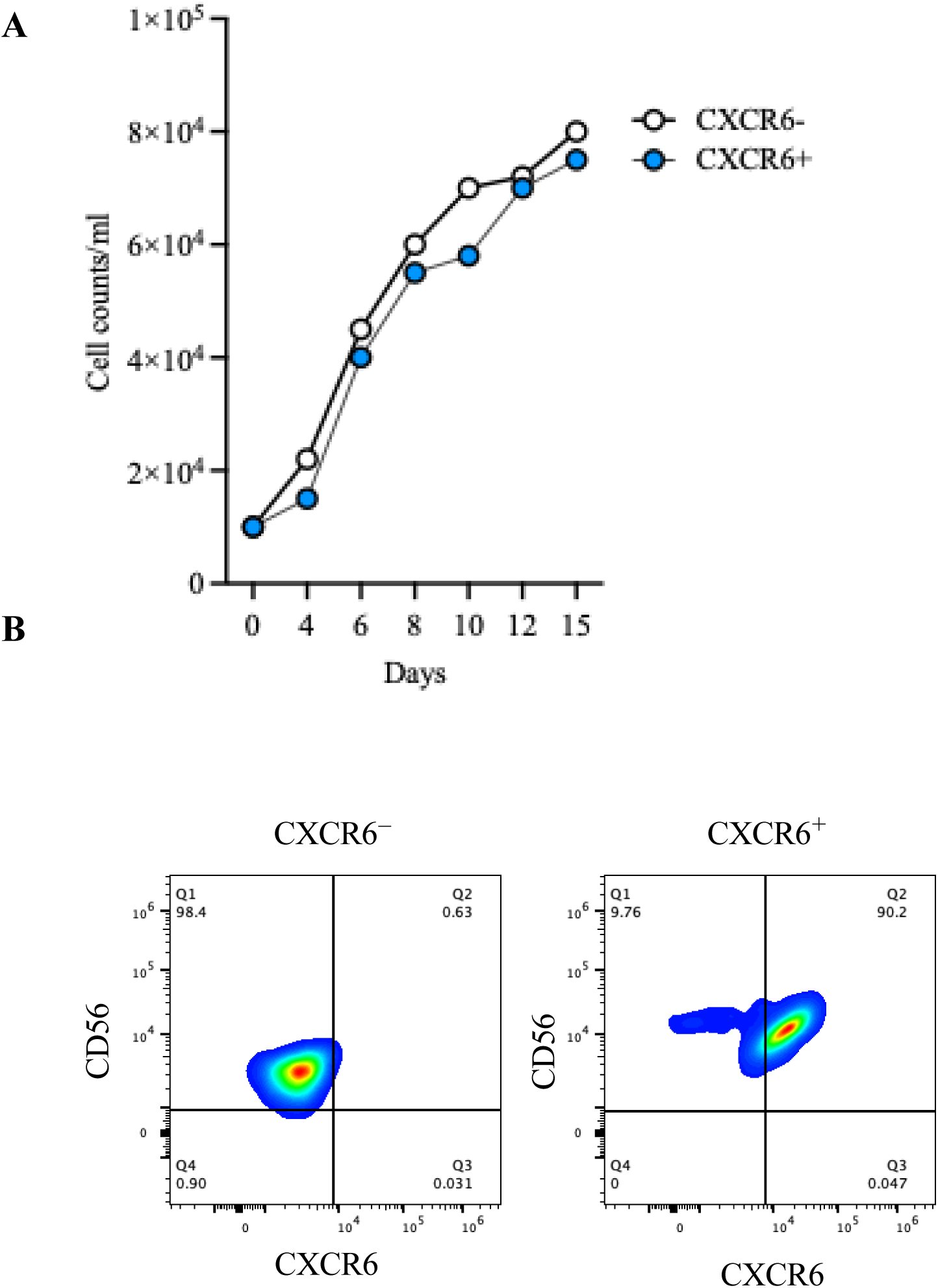
In vitro expansion and post-expansion purity of HIV-Env-vaccinated humanized mouse liver-derived CXCR6⁺ and CXCR6⁻ NK cells were assessed by cell counting and flow cytometry. **A)** Growth kinetics of expanded CXCR6⁺ and CXCR6⁻ NK cells are shown as cell counts per milliliter over 15 days. **B)** Representative flow cytometry plots show CD56 and CXCR6 expression in expanded CXCR6⁻ and CXCR6⁺ NK cell populations as determined by flow cytometry.

**Supplemental Figure 4.**
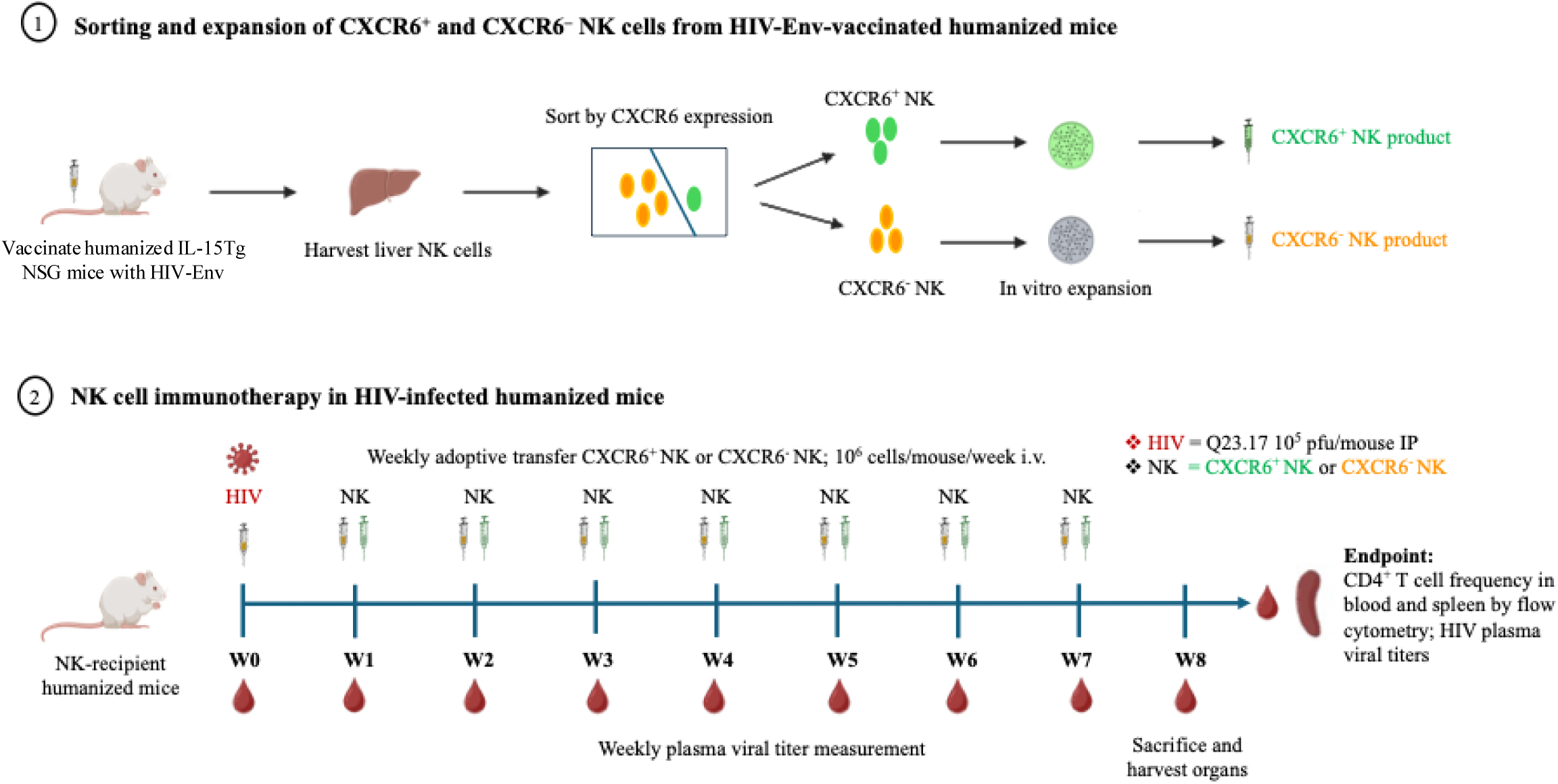
Experimental outline for NK cell immunotherapy using HIV Env-primed liver K cells. IL-15TgNSG mice were vaccinated with HIV Env, and liver NK cells were harvested, sorted based n CXCR6 expression, and expanded in vitro to generate CXCR6^+^ and CXCR6^−^ NK cell products. Umbilical ord blood-derived CD34^+^ HSC-humanized IL-15TgNSG mice were infected intraperitoneally with HIV-1 23.17, followed by weekly intravenous adoptive transfer of PBS control, 10⁶ CXCR6^+^ NK cells, or 10⁶ XCR6^−^ NK cells per mouse. Plasma was collected longitudinally each week to measure HIV viral burden, nd mice were sacrificed at week 8 for endpoint analysis of CD4^+^ T cell preservation in blood and spleen by low cytometry, together with assessment of plasma viral titers.

### 1 Supplementary Tables

**Supplementary table 1.** Flow cytometry Abs staining.

| Flowcytometry markers | Fluorophores | Clones |
| --- | --- | --- |
| hCD45 | BV605 | HI30 |
| mCD45 | AF532 | 30-F11 |
| CD56 | BV570 | HCD56 |
| DNAM-1 | BUV563 | DX11 |
| CD57 | PacBlue | HCD57 |
| CD16 | APC-Fire750 | 3GB |
| LILRB1 | PE-Cy7 | HP-F1 |
| NKp46 | BV650 | 9-E-2 |
| NKG2D | BUV737 | 1D11 |
| NKG2C | BUV615 | 134591 |
| 2B4 | BV421 | 2-69 |
| NKp30 | BUV805 | P30-15 |
| KIR3DL1 | PerCP-Cy5.5 | DX-9 |
| KIR2DL1 | AF488 | 143211 |
| KIR2DL3 | AF700 | 180701 |
| NKG2A | APC | Z199 |
| CXCR6 | PE/Dazzle 594 | K041E5 |
| CD69 | BUV496 | FN50 |
| CD107a | BUV395 | H4A3 |
| CD3 | APC/Fire | SK7 |
| Granzyme B | BV510 | GB11 |
| Perforin | BV711 | Dg9 |
| IFN-γ | BV750 | 4S.B3 |
| T-bet | BV786 | 4B10 |
| Ki67 | BV480 | B56 |
| EOMES | PerCp-eflour 710 | WD1928 |
| CXCR6 | BUV737 | RUO |
| mCD45 | Pacific Blue | 30-F11 |
| CD4 | BV750 | RU0 |
| HIV Gag p24 | FITC | Kc57-FITC |
| DAPI Live/Dead | Sigma Aldrich |  |

**Table S2.**
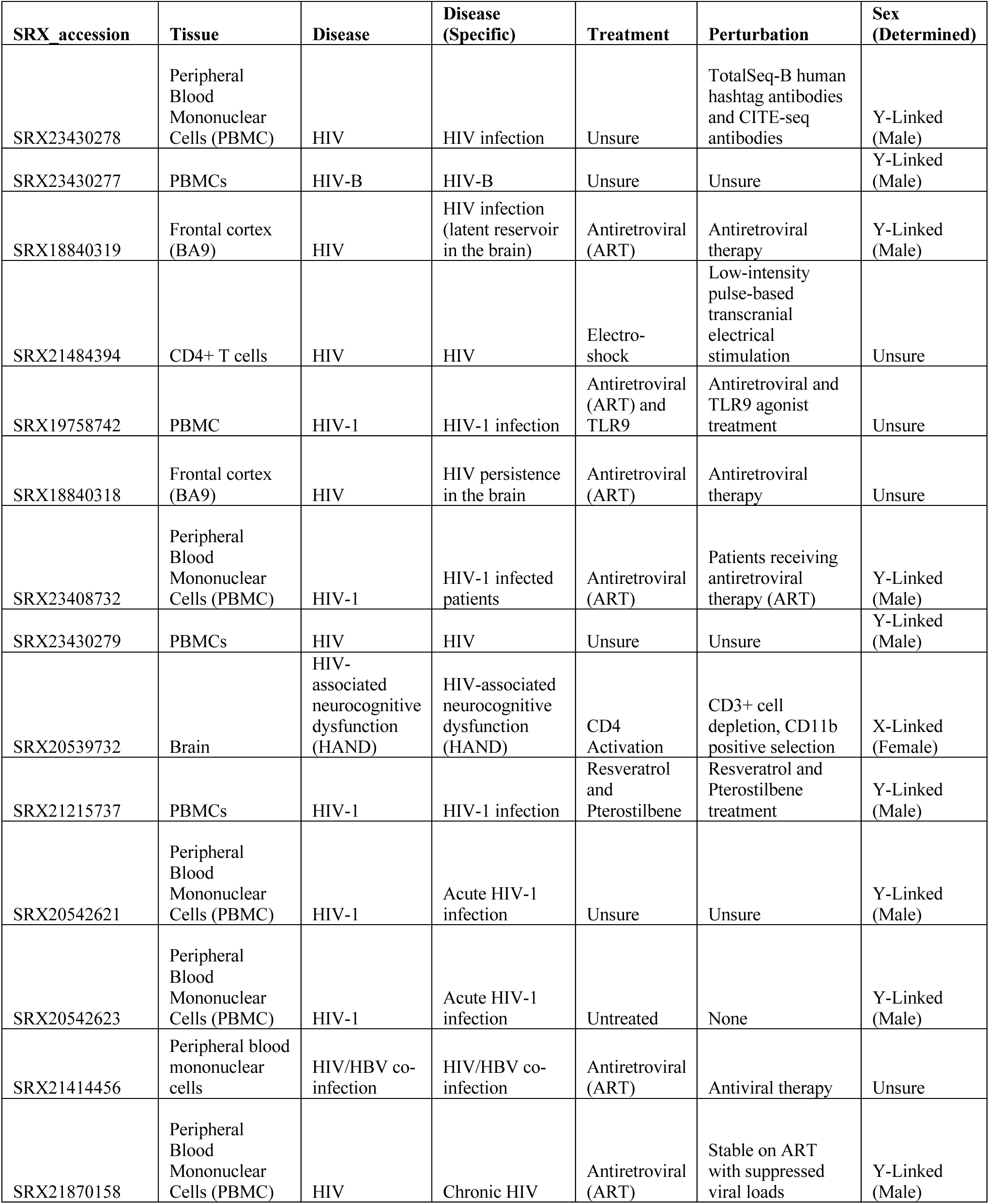
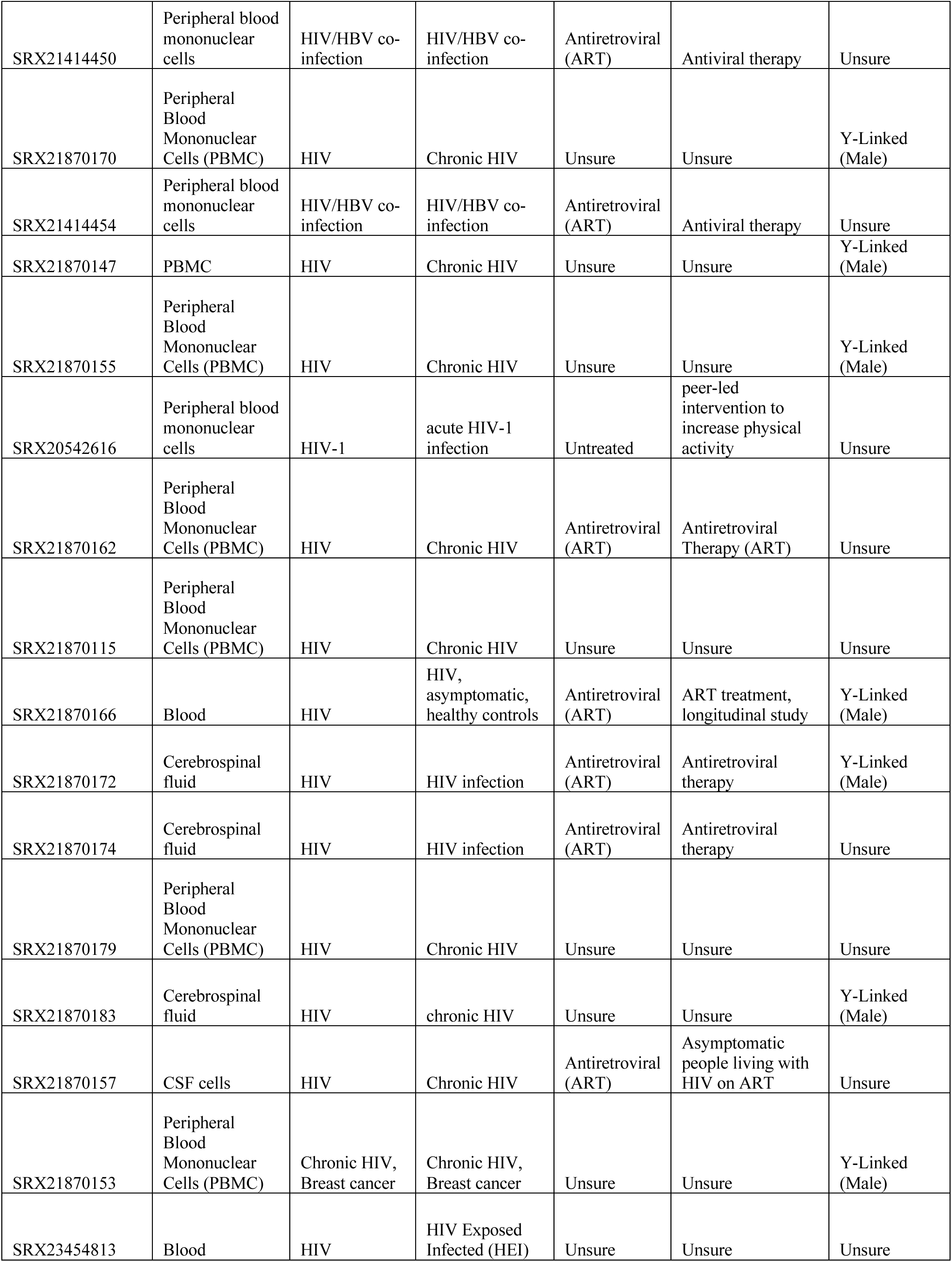
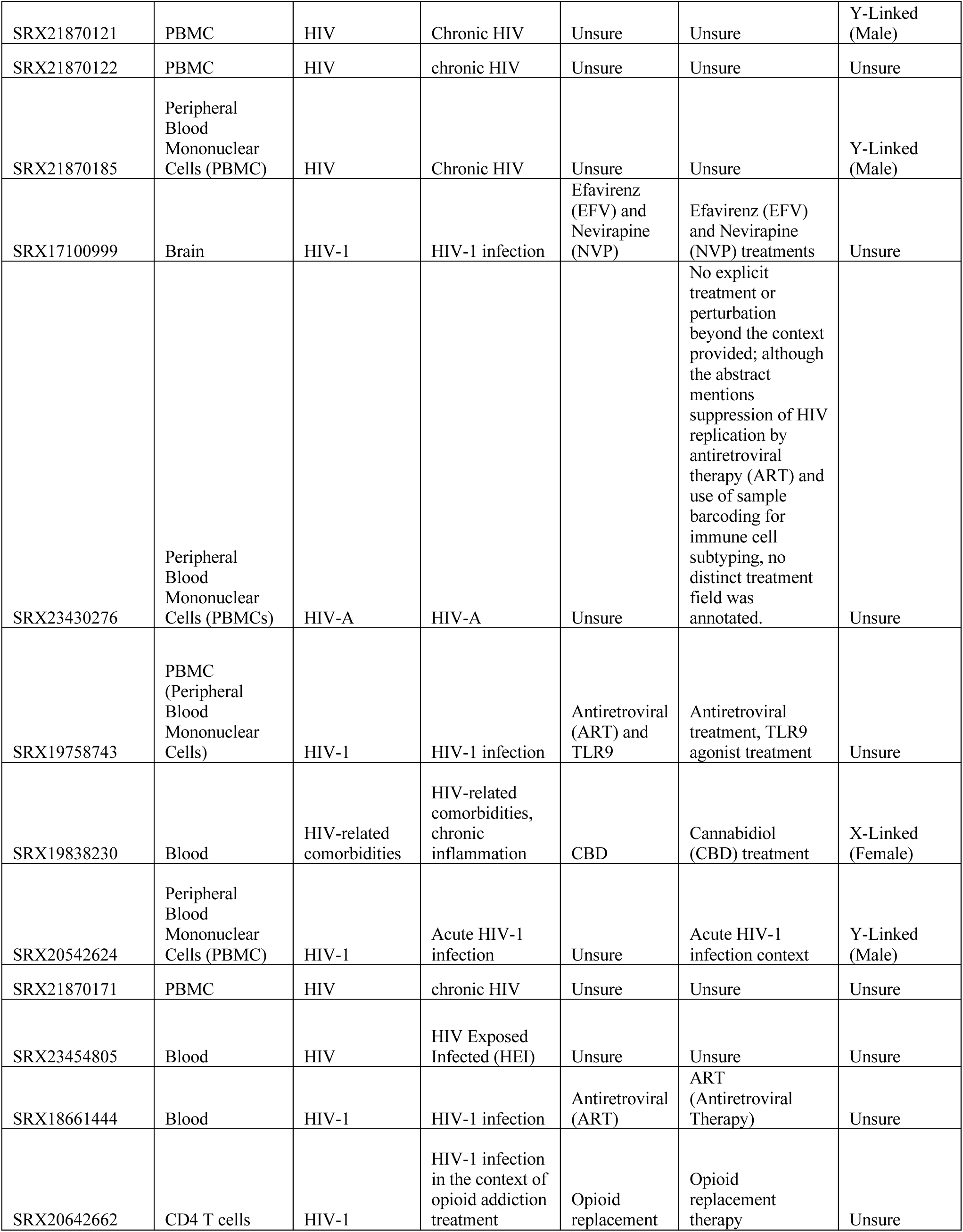
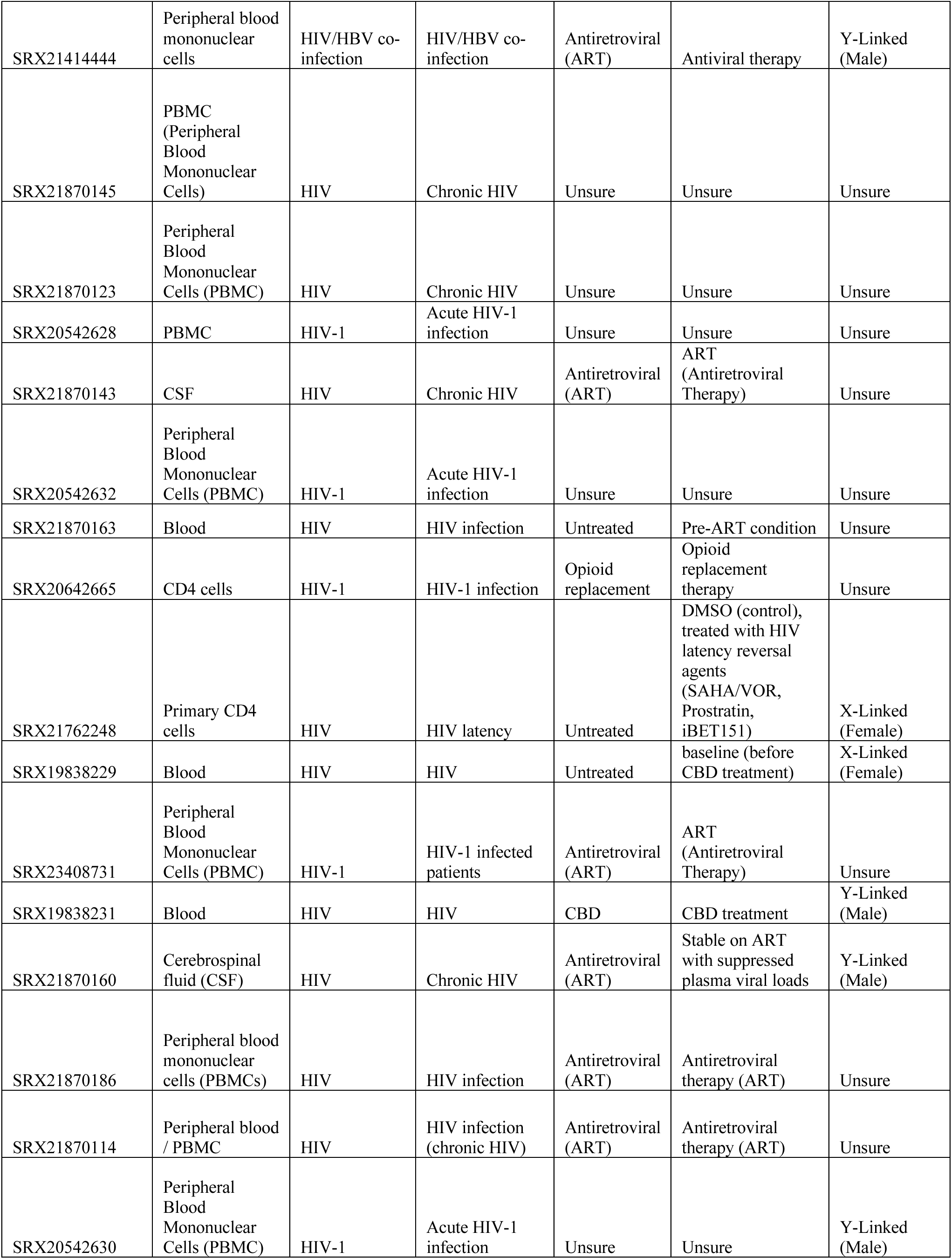
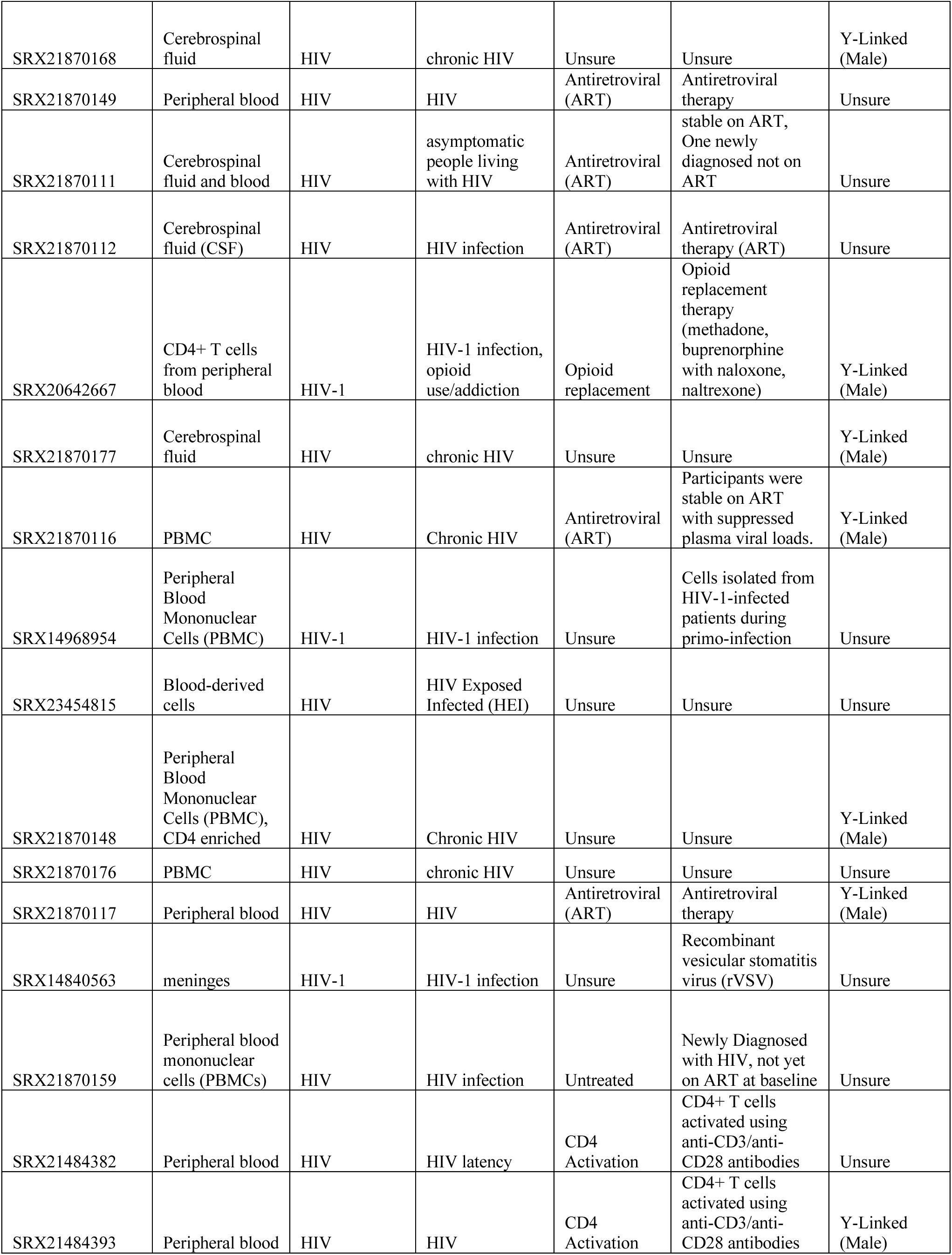
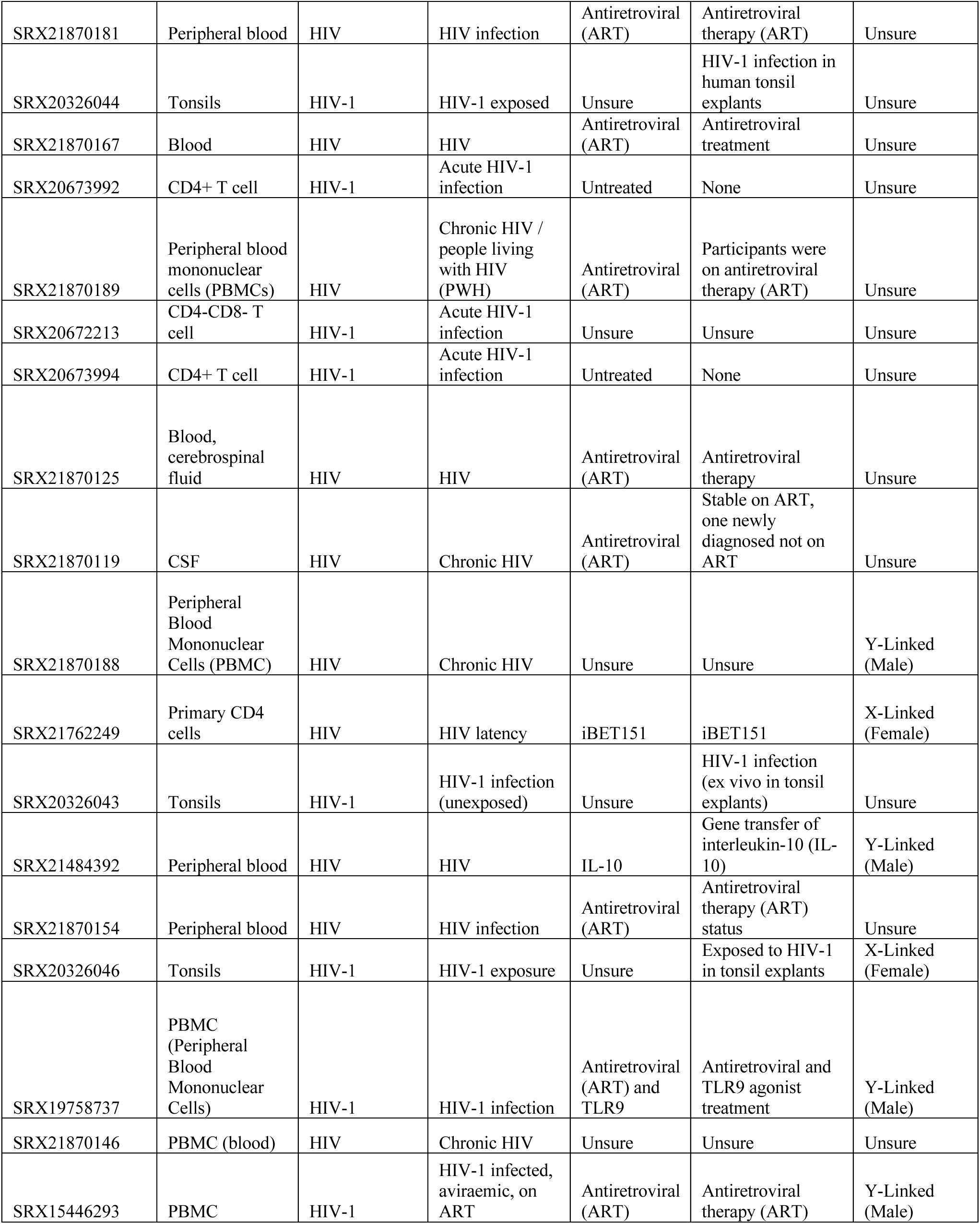
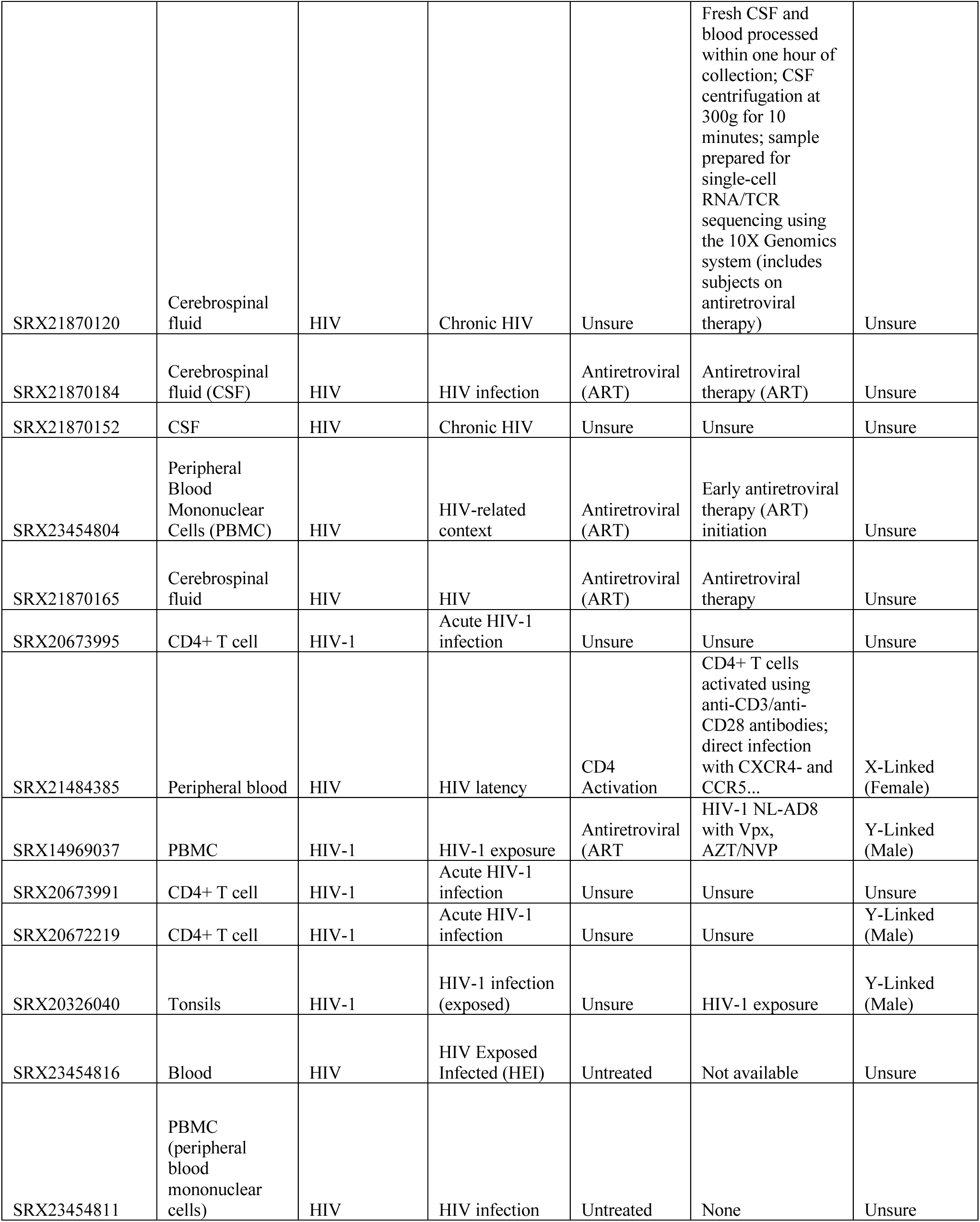
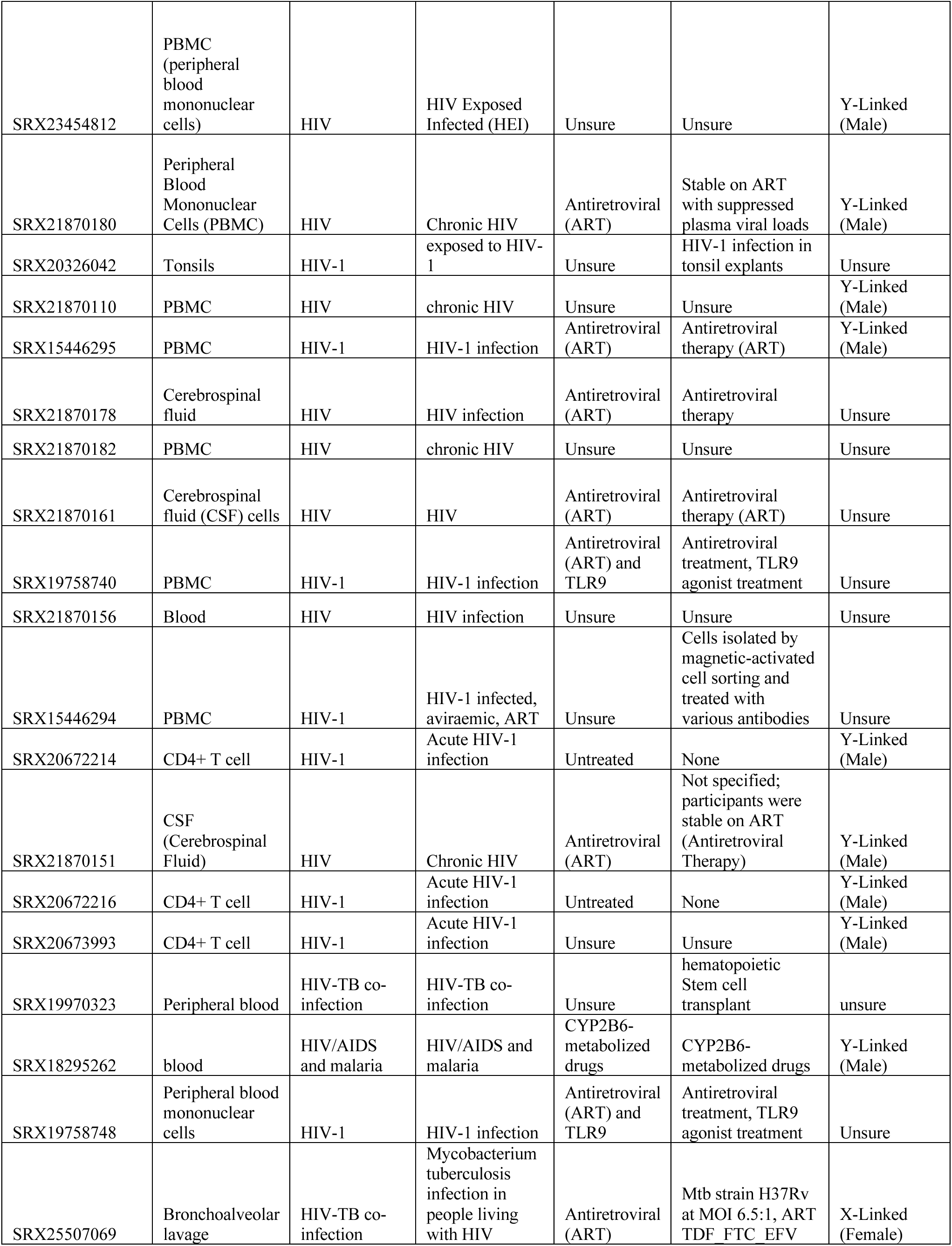
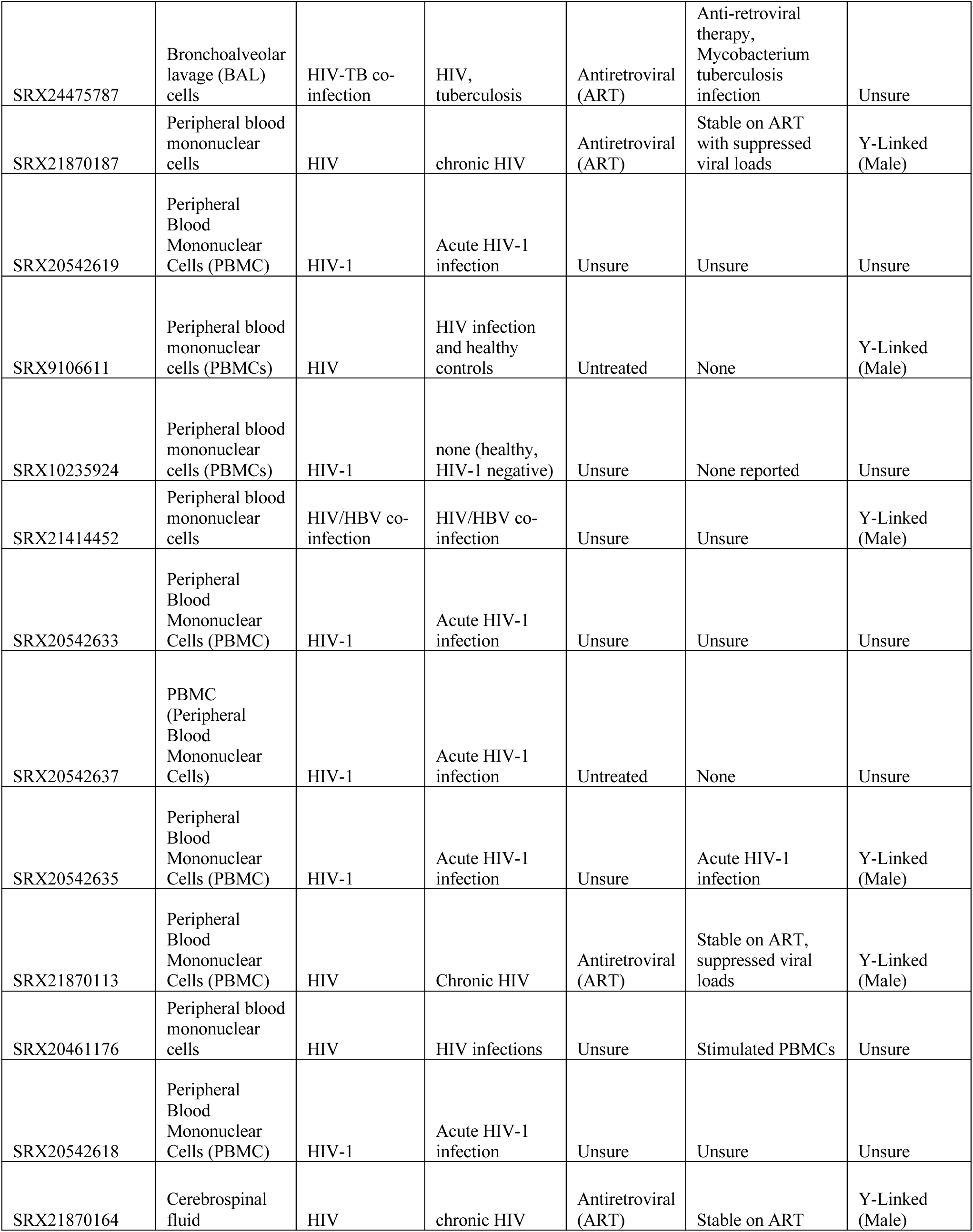
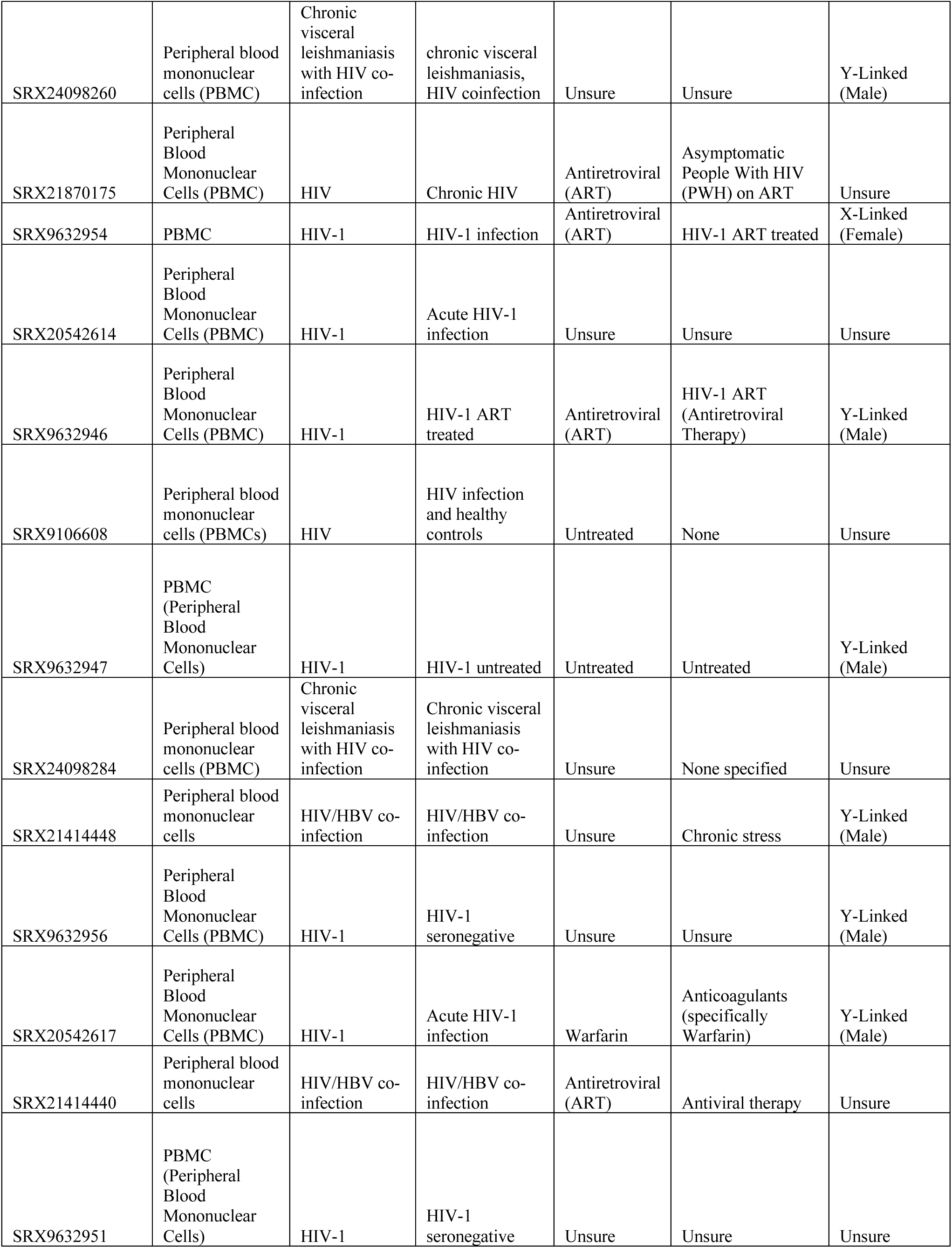
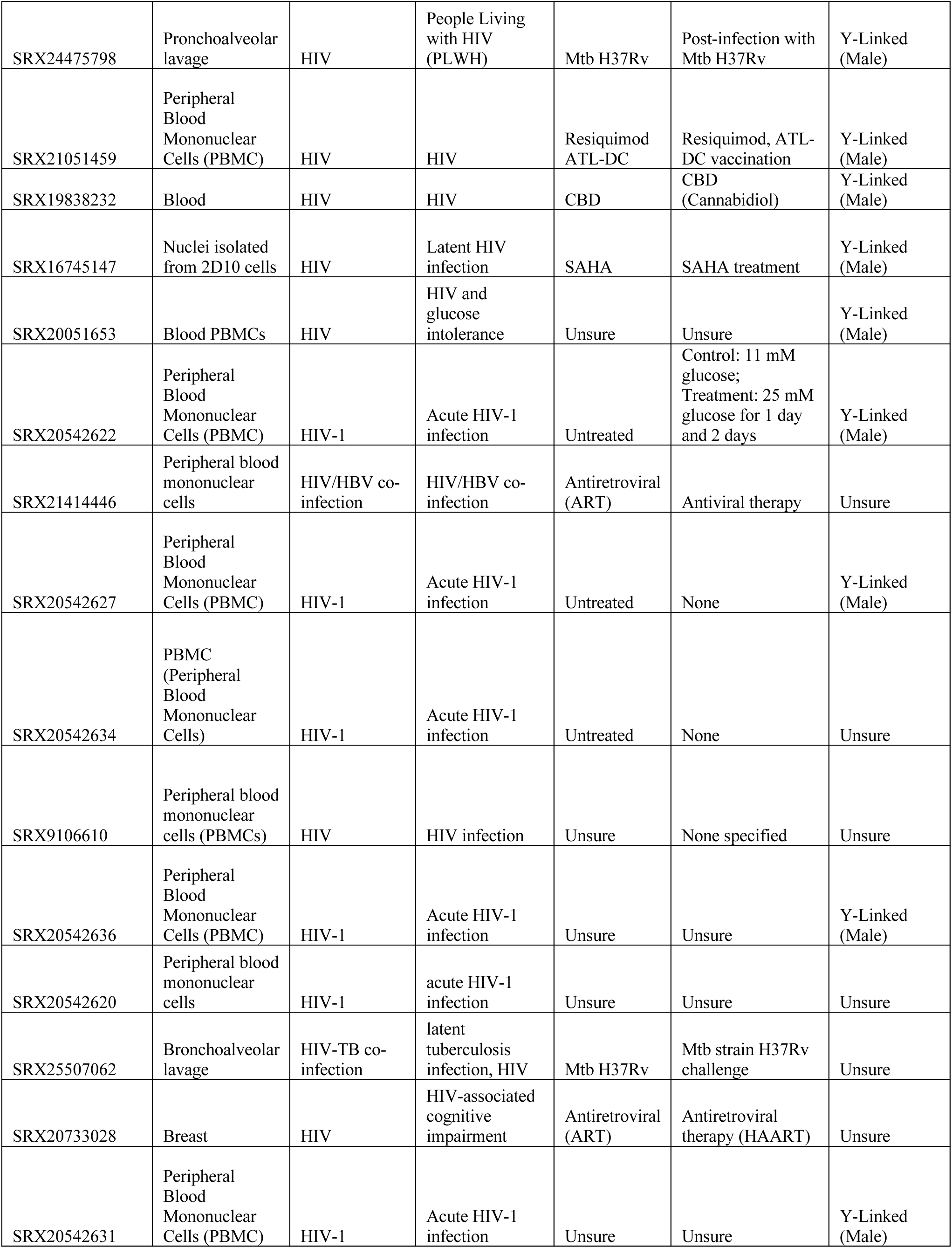
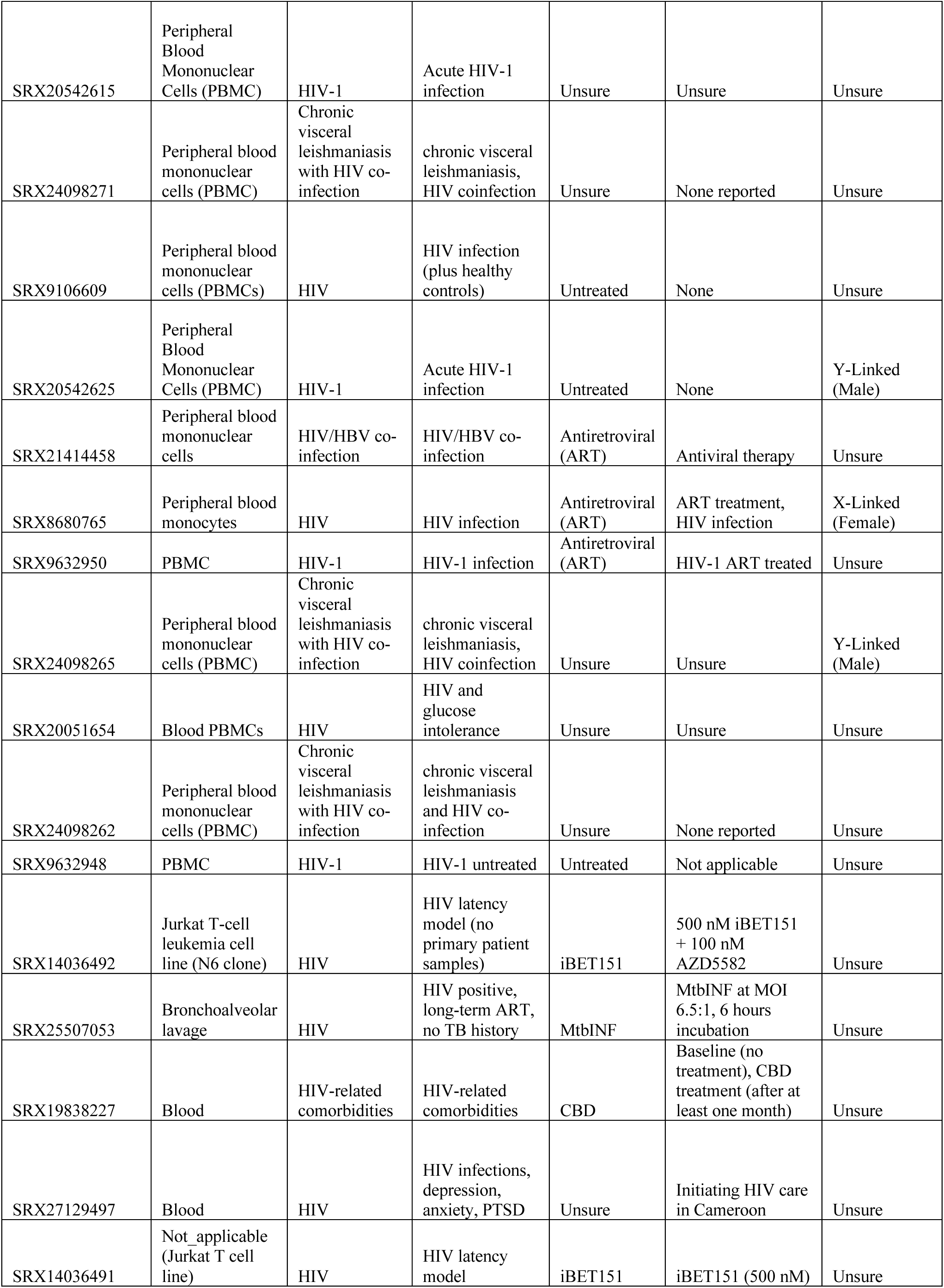
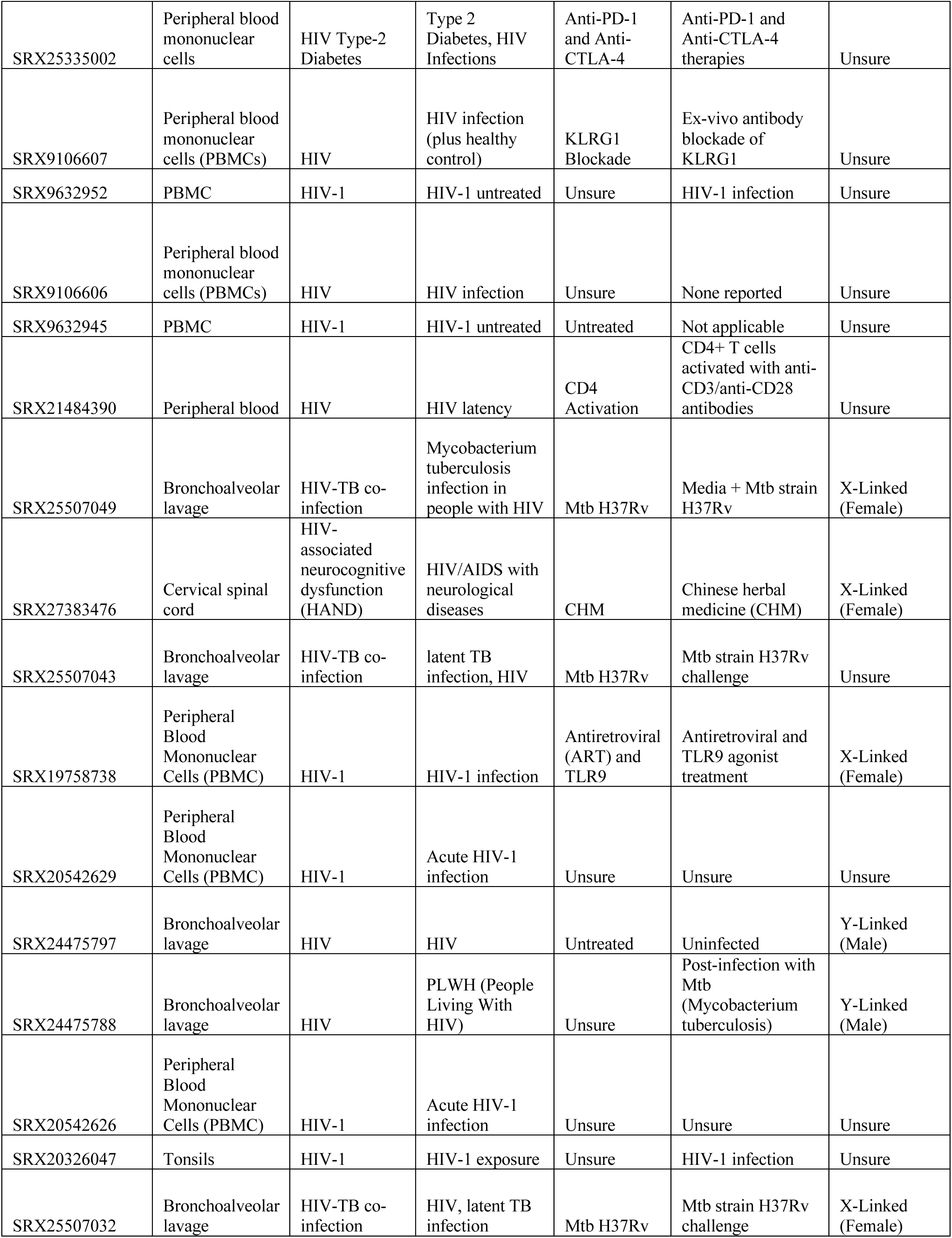
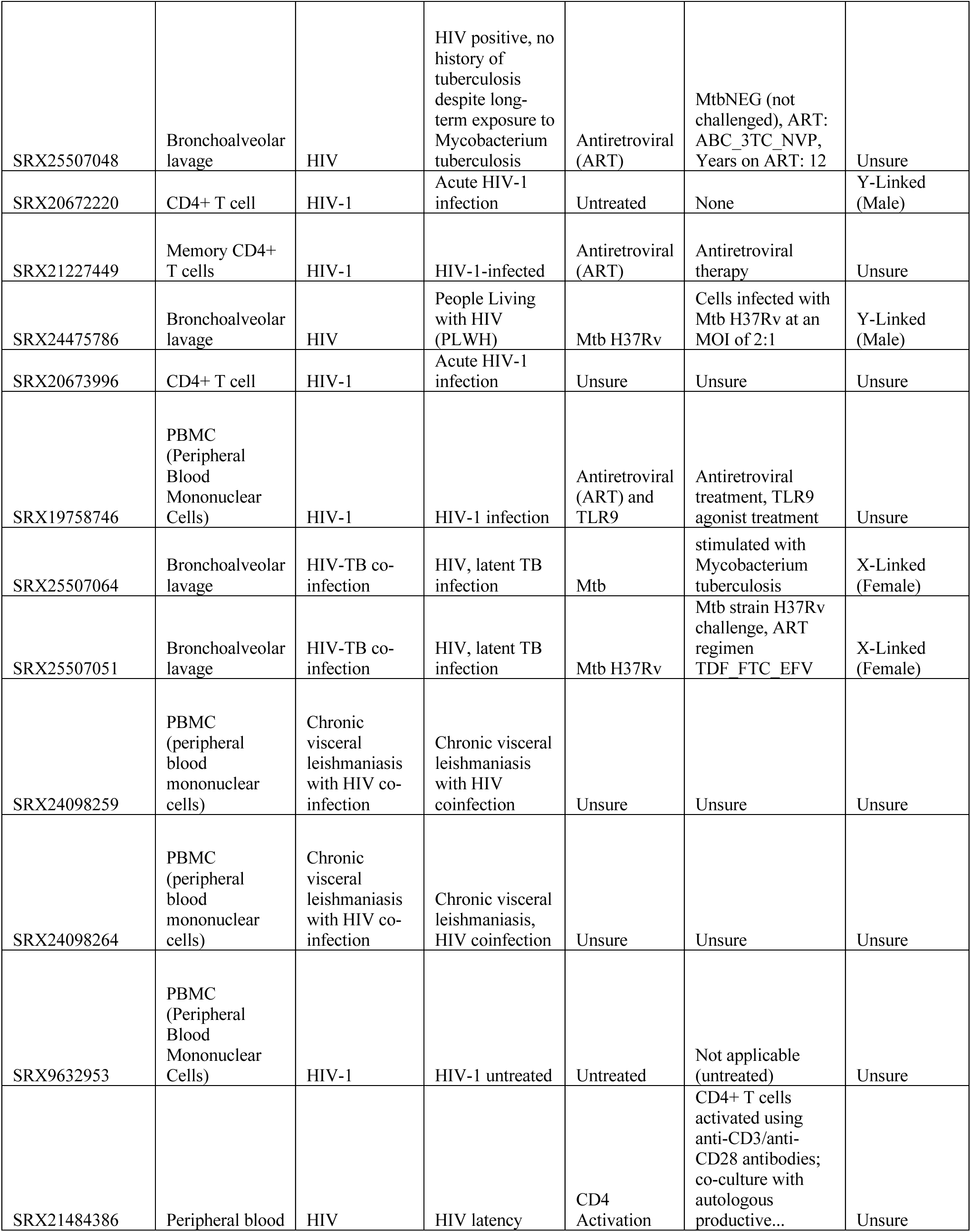
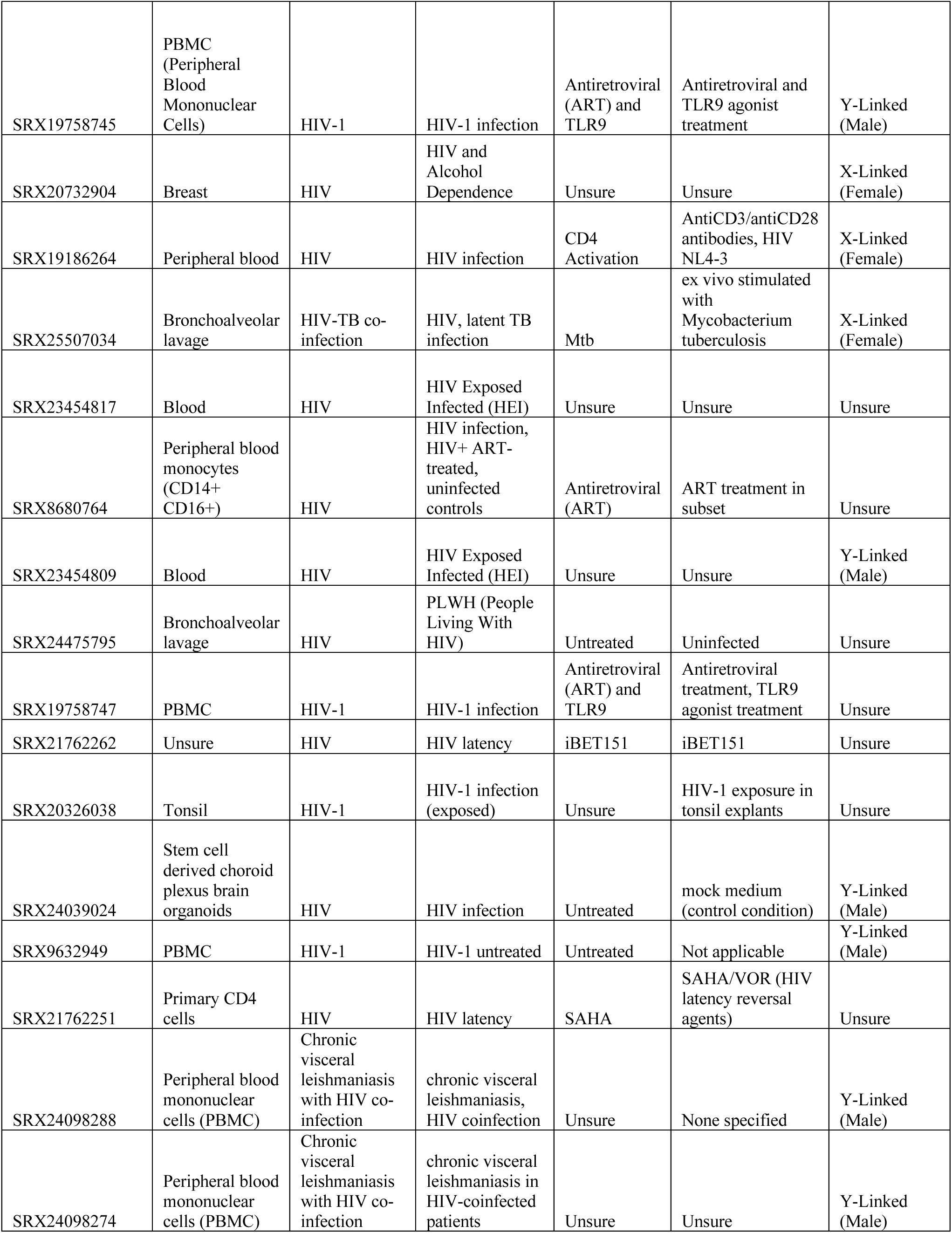
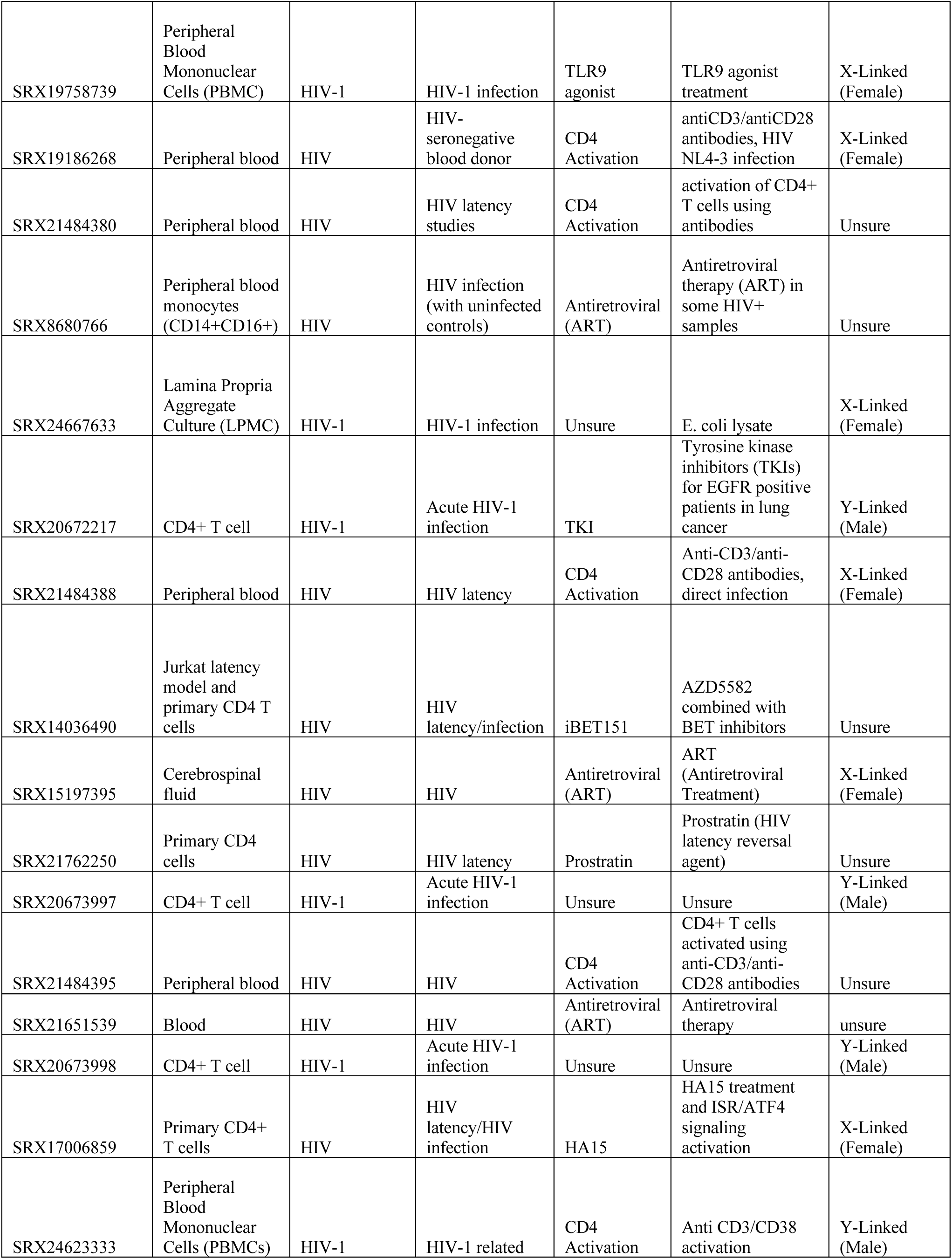
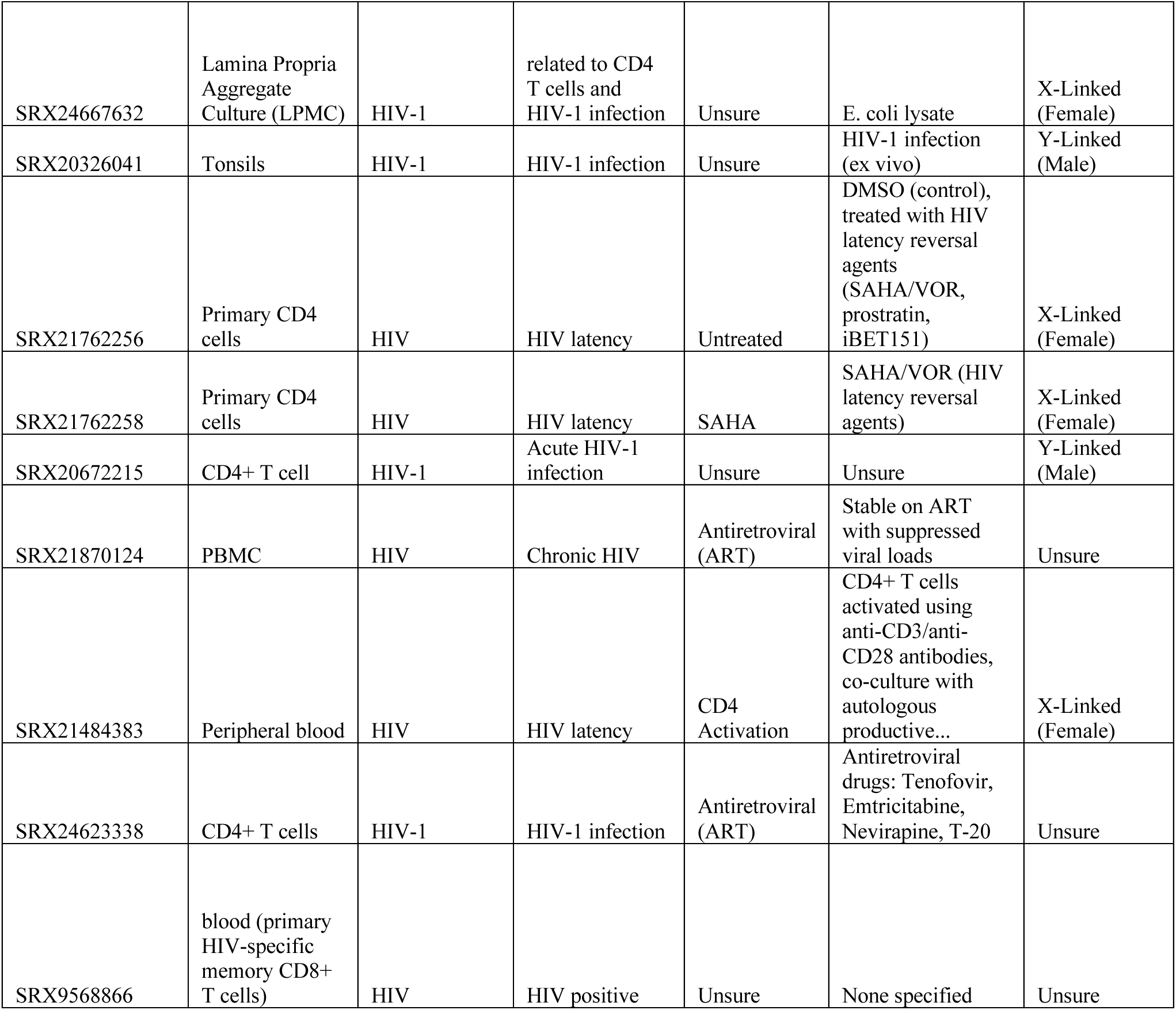
Source and Metadata for Figure 1.

**Supplementary Table 3.**
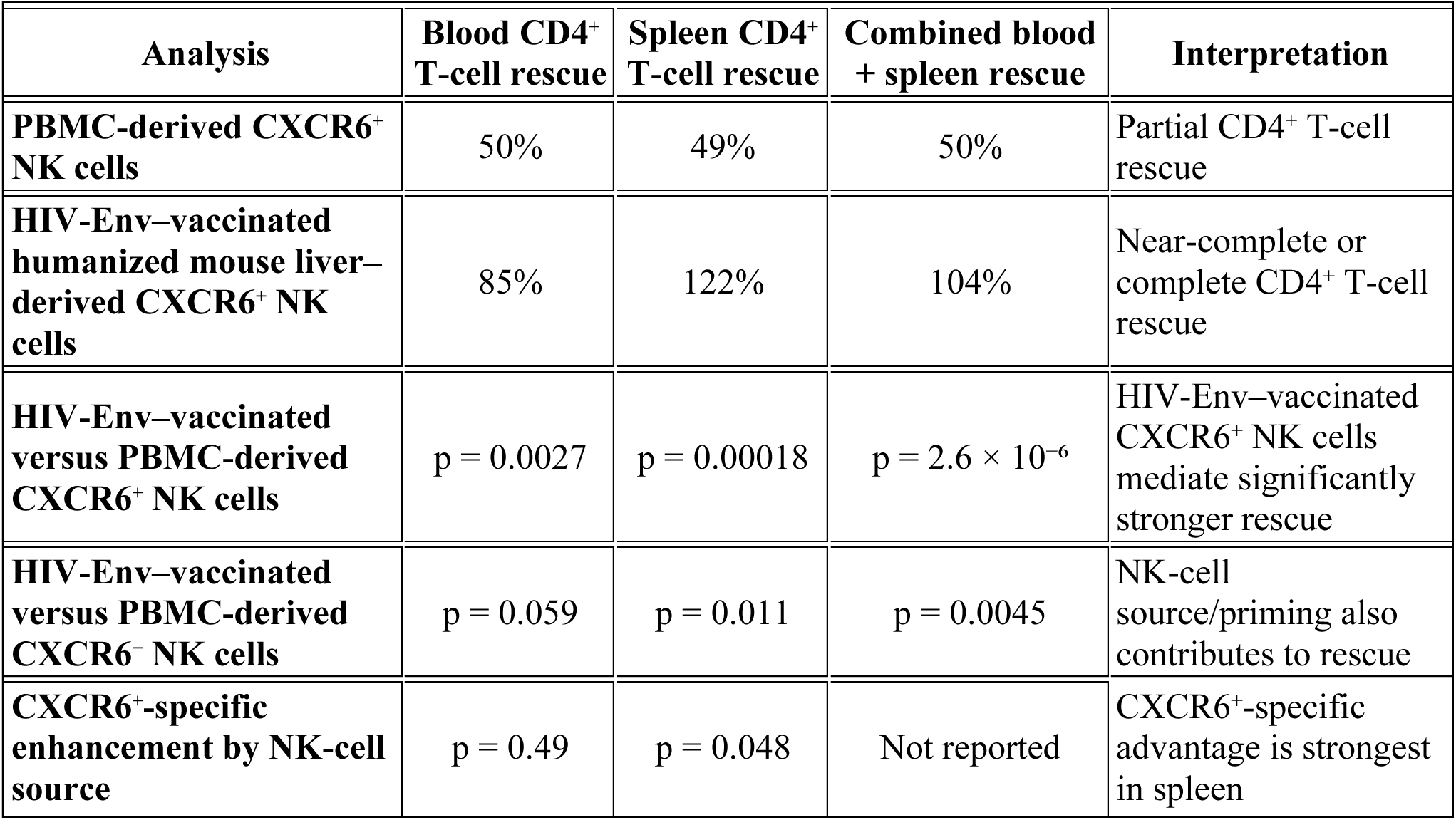
Normalized cross-experiment CD4⁺ T-cell rescue analysis comparing PBMC-derived and HIV-Env-vaccinated humanized mouse-derived NK-cell treatments.

